# Characterisation of protease activity during SARS-CoV-2 infection identifies novel viral cleavage sites and cellular targets with therapeutic potential

**DOI:** 10.1101/2020.09.16.297945

**Authors:** Bjoern Meyer, Jeanne Chiaravalli, Stacy Gellenoncourt, Philip Brownridge, Dominic P. Bryne, Leonard A. Daly, Arturas Grauslys, Marius Walter, Fabrice Agou, Lisa A. Chakrabarti, Charles S. Craik, Claire E. Eyers, Patrick A. Eyers, Yann Gambin, Andrew R. Jones, Emma Sierecki, Eric Verdin, Marco Vignuzzi, Edward Emmott

## Abstract

SARS-CoV-2 is the causative agent behind the COVID-19 pandemic, and responsible for over 170 million infections, and over 3.7 million deaths worldwide. Efforts to test, treat and vaccinate against this pathogen all benefit from an improved understanding of the basic biology of SARS-CoV-2. Both viral and cellular proteases play a crucial role in SARS-CoV-2 replication, and inhibitors targeting proteases have already shown success at inhibiting SARS-CoV-2 in cell culture models. Here, we study proteolytic cleavage of viral and cellular proteins in two cell line models of SARS-CoV-2 replication using mass spectrometry to identify protein neo-N-termini generated through protease activity. We identify previously unknown cleavage sites in multiple viral proteins, including major antigenic proteins S and N, which are the main targets for vaccine and antibody testing efforts. We discovered significant increases in cellular cleavage events consistent with cleavage by SARS-CoV-2 main protease, and identify 14 potential high-confidence substrates of the main and papain-like proteases, validating a subset with *in vitro* assays. We showed that siRNA depletion of these cellular proteins inhibits SARS-CoV-2 replication, and that drugs targeting two of these proteins: the tyrosine kinase SRC and Ser/Thr kinase MYLK, showed a dose-dependent reduction in SARS-CoV-2 titres. Overall, our study provides a powerful resource to understand proteolysis in the context of viral infection, and to inform the development of targeted strategies to inhibit SARS-CoV-2 and treat COVID-19.

## Introduction

SARS-CoV-2 emerged into the human population in late 2019, as the latest human coronavirus to cause severe disease following the emergence of SARS-CoV and MERS-CoV over the preceding decades (1, 2). Efforts to develop vaccines and therapeutic agents to treat COVID-19 are already yielding results, however it is widely expected that this first generation of treatments might provide imperfect protection from disease. As such, in-depth characterisation of the virus and its interactions with the host cell can inform current and next-generation efforts to test, treat and vaccinate against SARS-CoV-2. Past efforts in this area have included the proteome, phosphoproteome, ubiquitome and interactome of SARS-CoV-2 viral proteins and infected cells (3–9). Proteolytic cleavage plays a crucial role in the life cycle of SARS-CoV-2, and indeed most positive-sense RNA viruses. Inhibitors targeting both viral and cellular proteases have previously shown the ability to inhibit SARS-CoV-2 replication in cell culture models (10–13). Here we present a first unbiased study of proteolysis during SARS-CoV-2 infection, and its implications for viral antigens, as well as cellular proteins that may represent options for antiviral intervention. Proteolytic cleavage of the two coronavirus polyproteins generates the various viral proteins needed to form a replication complex, required for transcription and replication of the viral genome and subgenomic mRNAs. The key viral enzymes responsible are the papain-like (PLP, nsp3) and main proteases (Mpro, nsp5). Aside from cleaving viral substrates, these enzymes can also act on cellular proteins, modifying or neutralising substrate activity to benefit the virus. A recent study highlighted the ability of the viral proteases to cleave proteins involved in innate immune signaling including IRF3, NLRP12 and TAB1 (14). However, there has yet to be an unbiased study to identify novel substrates of the coronavirus proteases in the context of viral infection. The identification of such substrates can identify cellular enzymes or pathways required for efficient viral replication that may represent suitable targets for pharmaceutical repurposing and antiviral intervention for the treatment of COVID-19.

Viral proteins can also be the targets of cellular proteases, with the most prominent example for coronaviruses being the cleavage of the spike glycoprotein by the cellular proteases FURIN, TMPRSS2 and Cathepsins (10, 11, 15, 16), but the exact cleavage sites within spike for most of these individual cellular proteases are not yet characterised. Proteolytic processing can also be observed for other coronavirus proteins, for example, signal peptide cleavage of SARS-CoV ORF7A (17) and caspase cleavage of the nucleocapsid protein (18, 19). The spike glycoprotein especially, forms the key or sole component of the vaccines currently in use. For a functional immune response, it is vital that the antigens presented to the immune system, as part of these vaccines, closely mimic those seen in natural infection. An understanding of any modifications to these antigens observed during natural infection, such as glycosylation, phosphorylation and proteolytic cleavage, is critical to enable the rational design and validation of vaccine antigens and the selection of appropriate systems for their production. Currently a range of vaccine platforms are being explored, and certain platforms or delivery routes more likely to suffer from altered posttranslational modification states than others (20).

Mass spectrometry-based proteomic approaches have already led to rapid advances in our understanding of SARS-CoV2, with notable examples including the rapid release of the cellular interactome (6) and proximity interactome (7) for a majority of SARS-CoV-2 proteins, as well as proteomic (3, 5), phosphoproteomic (4, 8) and ubiquitomic analyses (9). Larger scale-initiatives have been launched focusing on community efforts to profile the immune response to infection, and provide in-depth characterization of viral antigens (21). Mass spectrometry has particular advantages for investigation of proteolytic cleavage as analysis can be conducted in an unbiased manner, and identify not only the substrate, but the precise site of proteolytic cleavage (22).

In this work we have applied mass spectrometry-based methods for N-terminomics to study proteolysis and the resulting proteolytic proteoforms generated in the context of SARS-CoV-2 infection, enabling the identification of novel cleavage and processing sites within viral proteins. We discovered several of these novel cleavage sites show altered cleavage following treatment with the cathepsin/calpain inhibitor calpeptin. We also identify cleavage sites within cellular proteins that match the coronavirus protease consensus sequences for Mpro and PLP, show temporal regulation during infection, are cleaved *in vitro* by recombinant Mpro and PLP, and demonstrate these proteins are required for efficient SARS-CoV-2 replication. These SARS-CoV-2 protease substrates include proteins that can be targeted with drugs in current clinical use to treat other conditions (23). Indeed, we demonstrate potent inhibition of SARS-CoV-2 replication with two compounds that are well-established chemical inhibitors of the SARS-CoV-2 protease substrates SRC and myosin light chain kinase (MYLK).

## Results

### Proteomic analysis of SARS-CoV-2-infected cell lines identifies alterations to the N-terminome

To investigate proteolysis during SARS-CoV-2 infection, N-terminomic analysis at various timepoints during the course of SARS-CoV-2 infected Vero E6 and A549-Ace2 cells (Fig. 1A) was performed. Vero E6 cells are an African Green Monkey kidney cell line commonly used for the study of a range of viruses, including SARS-CoV-2 which replicates in this cell line to high titres. A549-Ace2 cells are a human lung cell line which has been transduced to overexpress the ACE2 receptor to allow for SARS-CoV-2 entry. Cells were infected in biological triplicates at a multiplicity of infection (MOI) of 1, and harvested at 4 timepoints (0, 6, 12 and 24h) post-infection. Mock-infected samples were collected at 0h and 24h post-infection. These timepoints were chosen to cover SARS-CoV-2 infection from virus entry, over replication to virus egress: RNA levels increased from 9h post-infection (Fig. 1B), protein levels showed steady increases throughout infection (Fig. 1C), and viral titres increased at the 24h timepoint (Fig. 1D). These features were shared in both cell lines, with the Vero E6 cells showing greater RNA and protein levels, as well as viral titres compared with the A549-Ace2 cells. Analysis of the N-termini-enriched samples was performed by LC-MS/MS following basic reverse phase fractionation. For the purposes of this analysis, neo-N-termini were taken to be those beginning at amino acid 2 in a given protein or later. By this definition these neo-N-termini will include those with post-translational removal of methionine, signal peptide cleavage, as well as those cleaved by viral or cellular proteases. The modified N-terminomic enrichment strategy used (22) employed isobaric labelling (TMTpro) for quantification as this permitted all samples to be combined prior to enrichment, minimising sample variability. This strategy meant that only those peptides with a TMTpro-labelled N-terminus or lysine residue were quantified. As only un-blocked N-termini are labelled with undecanal, this approach results in the selective retention of undecanal-tagged tryptic peptides on C18 in acidified 40% ethanol, with N-terminal and neo-N-terminal peptides enriched in the unbound fraction (22).

**Fig. 1.**
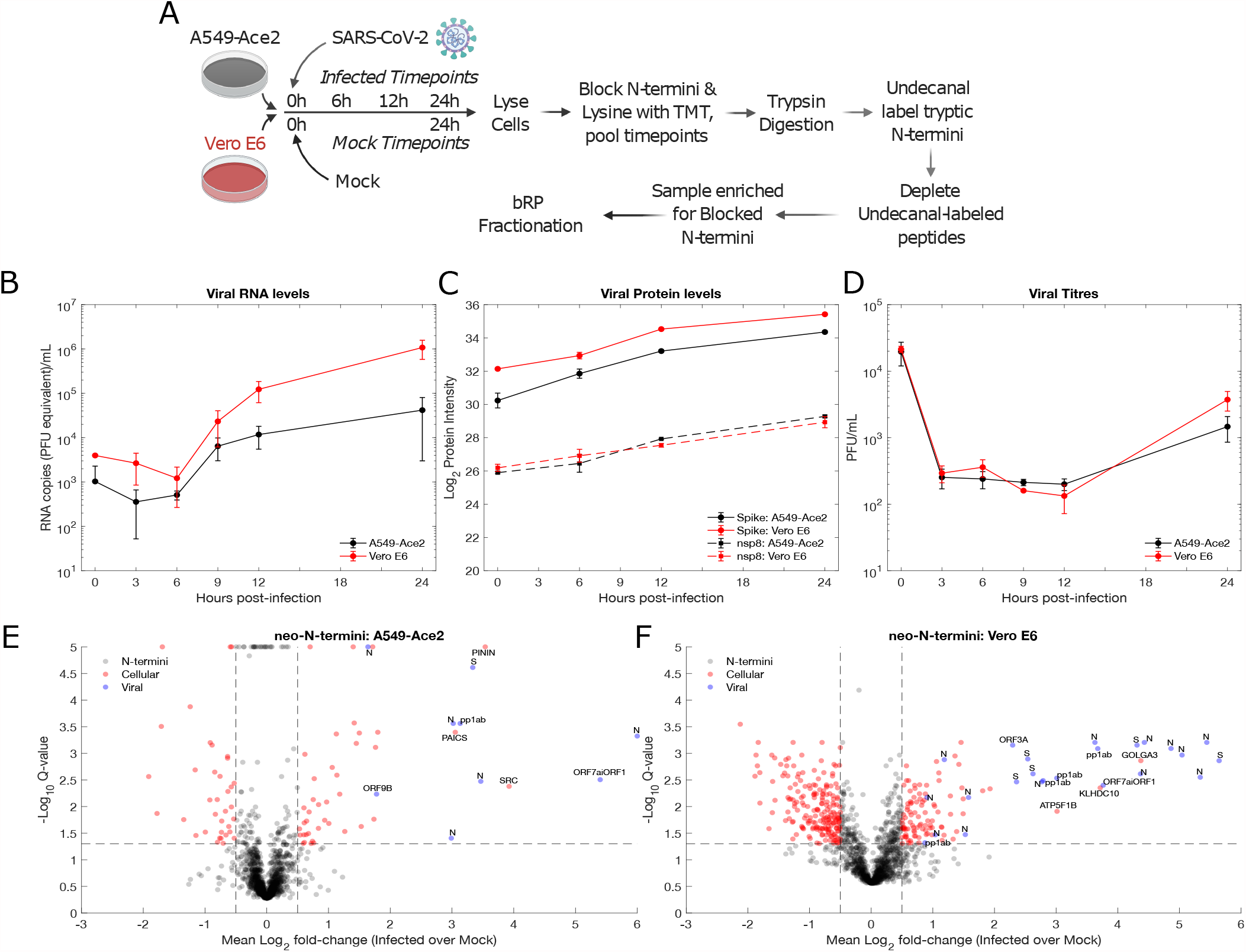
N-terminomic analysis of SARS-CoV-2 infection of A549-Ace2 and Vero E6 cells. A) Experimental design. B) Viral RNA levels were determined by qRT-PCR (n = 3 biological replicates). C) Protein levels were determined based on the TMTpro fractional intensity of the total protein intensity for the unenriched proteomic samples (n = 3 biological replicates). D) Infectious virus production (PFU). (n = 3 biological replicates). Error bars for B-D represent standard deviation. E) A549-Ace2 and F) Vero E6 neo-N-terminomic analysis reveals significant increases in peptides corresponding to viral and cellular neo-N-termini, where neo-N-termini must begin from amino acid 2 or later. P-values were obtained by two-tailed unpaired t-test, correction for multiple hypothesis testing to obtain Q-values was performed as described Storey (2002)(24). bRP = basic reverse phase fractionation, PFU = plaque-forming units. The vertical and horizontal lines correspond to fold change (0.5) cutoff and Q-value (0.05) cutoffs respectively.

Quality filtering of the dataset was performed (Fig. S1), infected and mock samples separated by PCA and 0h Mock, 0h infected and 6h infected clustered together, and away from the 12h and 24h infected samples (Fig. S1A-D). With the exception of the enriched Vero E6 dataset the 24h mock sample clustered with the 0h mocks. The Vero E6 24h Mock clustered away from the 0h and infected samples which may reflect regulation due to cell confluence as this was not observed with the paired unenriched sample. Sample preparation successfully enriched for blocked N-termini consisting of acetylated, pyroglutamate-N-termini and TMTpro-labelled N-termini (Fig. S1), and blocked N-termini were more abundant in the enriched samples. In both datasets, TMTpro-labelled N-termini represent approximately 50% of the blocked N-termini, with the rest split evenly between pyroglutamate and N-terminal acetylation (Fig. S1). After filtering, over 2700 TMTpro-labelled N-termini representing neo-N-termini were identified from each cell line. While the experimental design chosen is based around minimising missing data within rather than between the two datasets, 497 neo-N-termini were common to the two cell lines Fig. S2, and show positive correlation (Pearson’s *ρ*, 0.64-0.7).

When the 24h infected and mock-infected timepoints were compared, both cellular and viral neo-N-termini in A549-Ace2 (Fig. 1) and Vero E6 cells (Fig. 1F) were identified as showing significant alterations in their abundance. In line with expectation, N-termini from viral proteins were solely identified as showing increased abundance during infection in both cell lines. N-termini from cellular proteins showed both increased and decreased abundance during infection. We reasoned that those neo-N-termini showing increased abundance would include viral neo-N-termini, as well as those cellular proteins cleaved by the SARS-CoV-2 PLP and Mpro proteases. For this study we therefore focused specifically on viral N-termini and those cellular neo-N-termini identified as showing significantly increased abundance (t-test, multiple hypothesis testing corrected Q value*≤* 0.05) during infection.

### Novel proteolytic processing of SARS-CoV-2 proteins is observed during infection

The 30kb SARS-CoV-2 genome encodes a large number of proteins including two long polyproteins formed through ribosomal frameshifting, the structural proteins S, E, M and N and a range of accessory proteins (Fig. 2). Coronavirus proteins, in line with those of other positive-sense RNA viruses are known to undergo post-translational modifications, including proteolytic cleavage in some cases. Across all datasets we identified the S, M and N structural proteins, with the exception of E which has also not been observed in other proteomics datasets due to both short length and sequence composition (3, 5). We identified the ORF3a, ORF6, ORF8 and ORF9b accessory proteins, and all domains of the polyprotein aside from nsp6, 7 and 11.

**Fig. 2.**
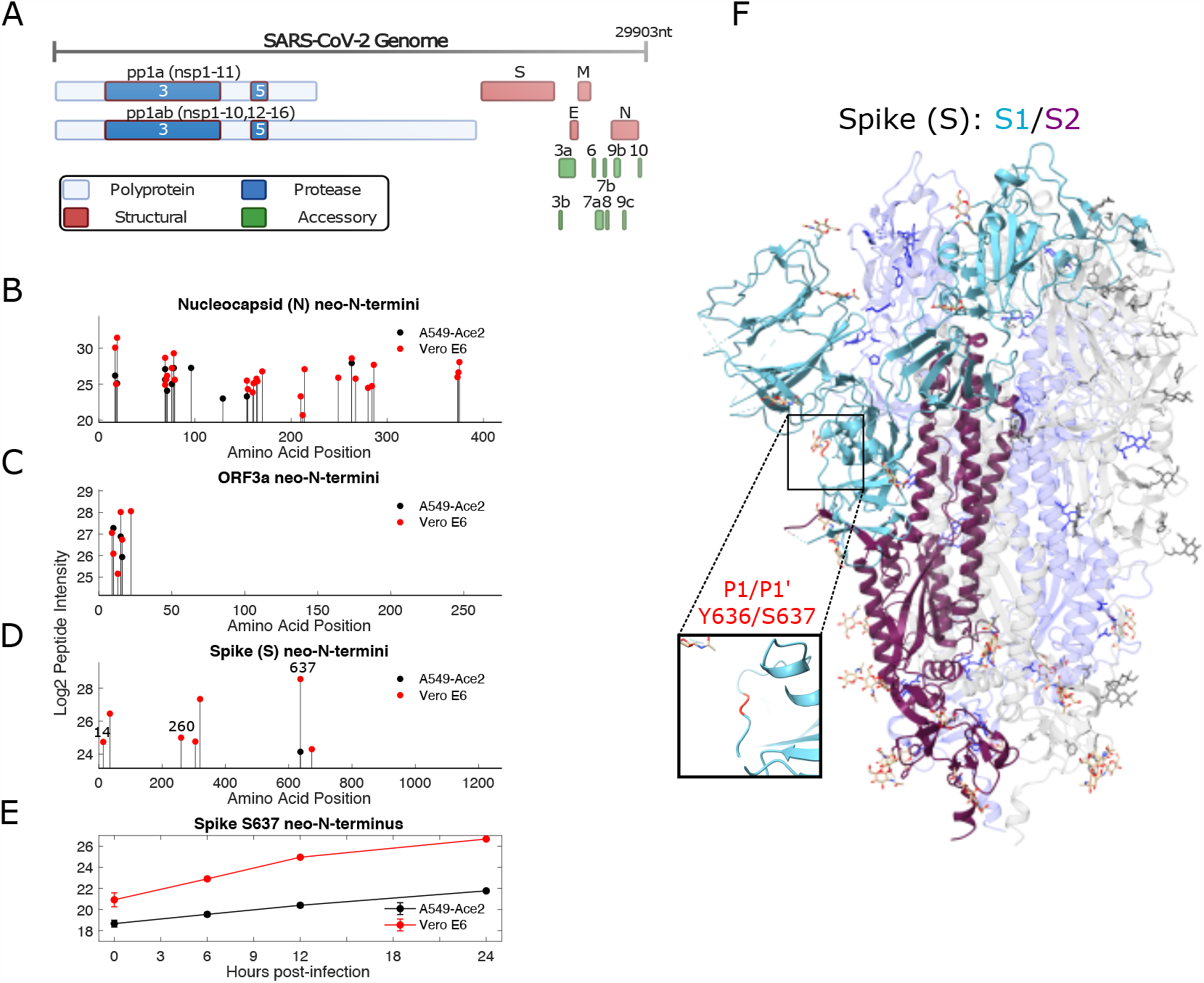
Proteolysis of viral proteins during SARS-CoV-2 infection. A) Schematic of the SARS-CoV-2 genome and proteome, with the nsp3 (PLP) and nsp3 (MPro) highlighted. Proteolytic processing of SARS-CoV-2 proteins during infection of A549-Ace2 and Vero E6 cells includes B) extensive cleavage of the nucleocapsid protein C) N-terminal processing of the ORF3a putative viroporin, and D) a novel cleavage site between Y636 and S637 in spike, N-terminal of the FURIN cleavage site. E) The abundance of the S637 spike neo-N-terminus increases over the infection timecourse (n = 3, error bars show standard deviation from 3 biological replicates). F) This cleavage site is present on an flexible region, C-terminal of the RBD (PDB: 6X6P).

We first sought to characterise neo-N-termini from viral proteins to understand potential patterns of cleavage that might generate functional proteolytic proteoforms of the viral proteins. Our search database included recently identified non-canonical translation products (25), and neo-N-termini and N-termini were identified from 8 viral proteins including the polyprotein (Fig. 2B-D; Fig. S3). Of these the nucleocapsid (N), ORF3a accessory protein and spike were most prominent. More cleavage sites were observed from infected Vero E6 cells than A549-Ace2 cells, which is in line with expectation given the higher levels of viral protein expression, and superior infectivity of this cell line compared to the A549-Ace2 cell line, permitting detection of less abundant cleavage products.

The coronavirus N protein is highly expressed during infection, and also represents a major antigen detected by the host immune response. Prior studies have identified cleavage of the SARS-CoV N protein by cellular proteases (18, 19), and our data identified multiple neo-N-termini consistent with proteolytic cleavage from both infected A549-Ace2 and Vero E6 cells (Fig. 2B). neo-N-termini common to both datasets include amino acids 17, 19, 69, 71, 76, 78, 154, and 263. Many of these cleavage sites were spaced closely together (e.g. 17/19, 69/71), consistent with a degree of further exoproteolytic processing. Some of these cleavage sites have subsequently been identified as autolysis products following extended incubations with N *in vitro* (26).

The ORF3a putative viroporin also shows N-terminal processing, possibly reflecting signal peptide cleavage (Fig. 2C). In a recent study, cryoEM of ORF3a in lipid nanodiscs did not resolve the first 39 N-terminal suggesting this region is unstructured (27). We observed N-terminal processing sites in the first 22 residues of the protein, with neo-N-termini beginning at amino acids, 10, 13 and 16 identified in both datasets, giving a possible explanation for the lack of N-terminal amino acids in cryoEM experiments.

Proteolytic cleavage of the spike glycoprotein is of major interest as it plays an important role in virus entry, with different distributions of cellular proteases between cell types resulting in the usage of different entry pathways, as well as potentially changing availability of surface epitopes for antibody recognition. Key proteases include FURIN, TM-PRSS2 and cathepsins, though in the latter two cases the actual cleavage sites targeted by these enzymes to process spike into S1 and S2 remain unclear. Consistent with previous observations (3, 5), we do not detect a neo-N-terminus deriving from the FURIN cleavage site as the trypsin digestion we employed would not be expected to yield peptides of suitable length for analysis. However, while beneficial for replication, FURIN cleavage is not essential and other cleavage events within spike can compensate (15, 16). We detect neoN-terminal peptides from S637 in both datasets (Fig. 2D). In line with the pattern of viral gene expression observed in the unenriched datasets this neo-N-terminus showed consistent increases in abundance throughout the experimental timecourse (Fig. 2E). S637 is located on a flexible loop near the FURIN cleavage site (Fig. 2F), suggesting it is accessible for protease cleavage (28). A mass spectrum for the S637 neo-N-terminus from the A549-Ace2 dataset is shown in Fig. S4A, the same peptide was observed with both 2+ and 3+ charge states in the Vero E6 dataset, and with a higher Andromeda score (124.37 vs. 104.82). Intriguingly, S637 was identified as a phosphorylation site in Davidson et al. (3). As phosphorylation can inhibit proteolytic cleavage when close the the cleavage site, this suggests potential post-translational regulation of this cleavage event.

Further neo-N-termini from spike were identified in the Vero E6 dataset alone, including a neo-N-terminus beginning at Q14 Fig. S4B. This is slightly C-terminal of the predicted signal peptide which covers the first 12 amino acids. This peptide featured N-terminal pyroglutamic acid formed by cyclization of the N-terminal glutamine residue. The peptide does not follow an R or K residue in the spike amino acid sequence and thus represents non-tryptic cleavage. The absence of TMTpro labelling at the N-terminus suggests that this N-terminus was blocked prior to tryptic digestion, with this modified N-terminus preventing TMTpro modification. Artifactual cyclization of N-terminal glutamine or glutamic acid residues typically results from extended trypsin digestion and acidic conditions (29). However, the order of labelling and digestion steps in our protocol, and non-tryptic nature of this peptide suggests that this N-terminal pyroglutamic acid residue is an accurate reflection of the state of this neo-N-terminus in the original biological sample. While this is to our knowledge, the first observation of this N-terminal modification and signal peptide cleavage site for spike during infection, cleavage and cyclization of Q14 has been observed with recombinantly-produced SARS-CoV-2 spike in HEK293 (30) and CHO cells (31). Three further N-terminal pyroglutamic acid residues were identified in SARS-CoV-2 proteins (N, ORF7A) within the Vero E6 dataset and can be found in table S2.

We detected viral neo-N-termini and N-termini in M, ORF7a, ORF9b and pp1ab. Due to conservation with SARS-CoV ORF7a, the first 15 residues of SARS-CoV-2 ORF7a are expected to function as a signal peptide which is posttranslationally cleaved (17, 32). neo-N-termini were identified in both datasets consistent with this hypothesis. Due to inclusion of the ORF7a iORF1 proposed N-terminal truncation of ORF7a which lacks the first two amino acids in ORF7a in the SARS-CoV-2 sequences used for data analysis, the start position of this neo-N-terminal peptide is given as 14 (25). However, this would be position 16 in ORF7a, consistent with removal of the signal peptide (MKIILFLAL-ITLATC, in Uniprot P0DTC7), and conserved with that in SARS-CoV ORF7a.

The native N-terminus of ORF9b was also identified, and several sites mapping to the replicase polyprotein, including a conserved neo-N-terminus consistent with predicted nsp10-nsp12 cleavage by Mpro. A neo-N-terminus consistent with nsp15-nsp16 cleavage by Mpro was identified in A549-Ace2 cells, and several internal neo-N-termini deriving from nsp1, -2 and -3 were also observed, though not common to both datasets. All the viral neo-N-termini and N-termini identified in this study can be found in tables S1 (A549-Ace2) and S2 (Vero E6) respectively. Table S3 includes all viral peptides identified in this study in both enriched and unenriched datasets.

### neo-N-termini and current Variants of Concern (VOC’s)

Currently there is extensive interest in mutations present in emerging Variants Of Concern (VOCs) and Variants of Interest (VOIs), and using our understanding of virus biology to predict how such mutations could alter protein function or antibody evasion.

To investigate if the viral neo-N-termini we identified may be under selective pressure, mutations characteristic of current VOC/VOIs were identified from covariants.org (33) on the 24th May 2021. These included B.1.1.7, B.1.351, B.1.427/9, B.1.525, B.1.526, P.1, B.1.617.1 and B.1.617.2. Compared to the a background proteome of all viral peptides identified in the analysis, neo-N-termini were not overrepresented near characteristic variant mutations (KS test, p = 0.146) Fig. S6. However, not all neo-N-termini may be under selective pressure, and in some cases mutations could be lethal to the virus so would not be observed in variants. While our data does not suggest global enrichment of neo-N-termini proximal to characteristic variant mutations, 9 of our neo-N-termini were within 5 amino acids of a characteristic variant mutation (table S6), and 17 were within 10 amino acids (table S7). Those neo-N-termini within 5 amino acids of variant mutations are of particular interest as the variant mutations could readily alter proteolytic cleavage. These variant mutations include multiple sites in S (13, 18, 19, 677), N (12, 80, 205, 377) and ORF3a (26). Of particular interest is the S13I mutation in spike in strain B.1.427/B.1.429, immediately preceding the pyroglutamate-modified Q14 neo-N-terminus. Recent research has indicated that this is tolerated in spite of blocking signal peptide cleavage to generate the Q14 neo-N-terminus, with signal peptide cleavage instead generating a neo-N-terminus at V16 resulting in loss of antibody binding and structural rearrangements within spike due to loss of disulphide bond formation between C15 and C136 (34). As further variants emerge, the incorporation of post-translational modification data from studies such as this can support efforts to predict phenotypes from genetic data on emerging variants.

### Novel SARS-CoV-2 cleavage sites are sensitive to calpeptin and a mutation proximal to the 637 cleavage site results in a higher fraction of cleaved spike in purified pseudovirus and enhanced cell entry

The N-terminomics experiments above successfully identified multiple previously uncharacterised proteolytic cleavage sites within viral proteins. However, we have limited information on the identity of the causal proteases behind these cleavage events. To address this, we performed a further N-terminomics experiment comparing the relative abundance of these cleavage sites and viral proteins following treatment with specific protease inhibitors Fig. 3A. The first, camostat mesylate is currently in clinical trials to treat COVID-19 disease and acts on TMPRSS2 and trypsin. It should be noted that the cell lines our study focuses on are TMPRSS2 negative and so camostat mesylate was included as a control. The second, calpeptin inhibits cathepsin and calpain cleavage. The experiment was performed in Vero E6 cells as the majority of viral cleavage sites identified in the first dataset were found in this cell line. Inhibitors were added at 12h post-infection with the aim of reducing proteolytic cleavage rather than inhibiting virus replication *per se* by permitting viral replication to proceed unimpeeded for the first 12h. Samples were then harvested at 24h post-infection Fig. 3A. Quality control of this dataset against showed tight clustering of the relevent samples, with infected samples clustering away from the mock-infected cells Fig. S5A,B. This dataset identified fewer quantifiable N-termini Fig. S5C-F, but these included the key cleavage sites in S and N.

**Fig. 3.**
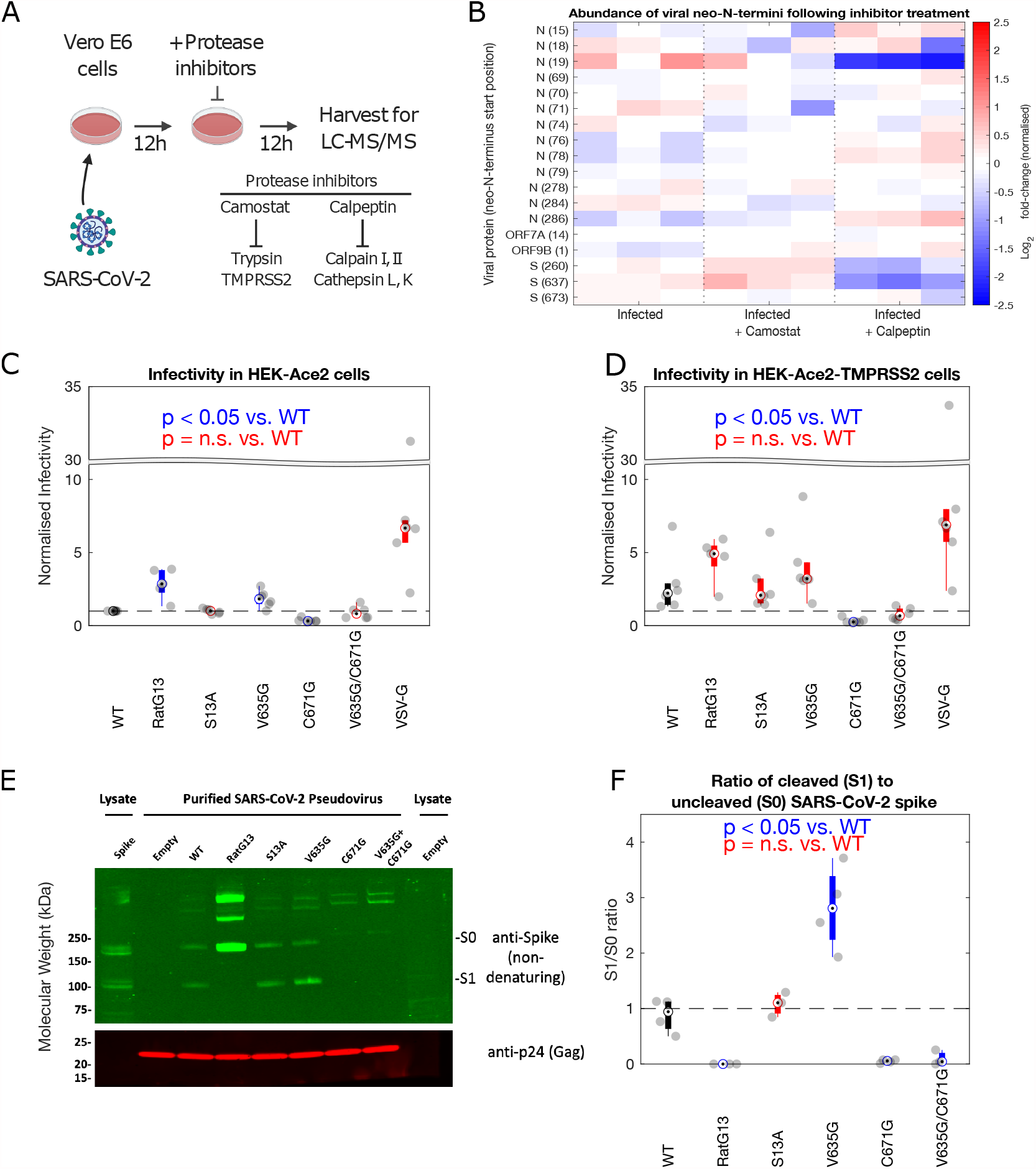
Several viral neo-N-termini show sensitivity to specific protease inhibitors and a spike 637-proximal mutation alters viral entry in TMPRSS2-ve cells. a) Experimental design for N-terminomics of SARS-CoV-2 infection in the presence of protease inhibitors. b) Abundance of viral neo-N-termini in infected cells +/-inhibitors. Data normalised to total levels of the relevant viral protein (n = 3 biological replicates). Pseudovirus entry assay conducted in c) HEK-Ace2 and d) HEK-Ace2-TMPRSS2 cells. The infectivity of lentivectors (LV) pseudotyped with the different spike mutants was normalized to that of WT in the HEK-Ace2 cell line (n *≥* 5 biological replicates) E) Western blotting of the pseudovirus stocks used in C) and D) confirms spike expression and incorporation into lentiviral particles. F) Densitometry analysis of spike western blotting data, examining the ratio between uncleaved (S0) and cleaved (S1) portions of the spike protein present in purified pseudotyped lentivirus stocks (n *≥* 3 biological replicates). Boxplots show median and interquartile ranges with whiskers extending to the most extreme non-outlier values. Unpaired Welch’s t-tests, which do not assume equal variance were used for statistical analyses.

Analysis of viral protein levels showed that when added at this late time post-infection viral protein levels were simiilar but did show some differences with the untreated infected samples Fig. S5G. ORF9B showed significantly reduced abundance in protease inhibitor-treated infected cells compared to infected but untreated cells (Camostat: t-test, p = 0.0031; Calpeptin: t-test, p = 0.0012). ORF3A showed significantly increased abundance compared to untreated-infected cells in the camostat-treated cells alone (t-test, p = 0.0016). Neither N or S showed significant changes in their abundance with either inhibitor treatment.

Neo-N-termini corresponding to viral proteins were normalised to the abundance of the viral protein from which the neo-N-terminus was derived and can be seen in Fig. 3B. The largest changes were observed following calpeptin treatment, resulting in significantly reduced abundance of neo-N-termini beginning at N(19),S(260) and S(637). Both neoN-termini from S (260, 637) showed increased abundance following treatment with camostat compared to untreated infected cells (t-test, p = 0.0074 and 0.0288 respectively). Two neo-N-termini within N (78, 286) show increased abundance in the calpeptin-treated infected cells (t-test, p = 0.0045, 0.0120). Reduced abundance of neo-N-termini following calpeptin inhibibion (e.g. N(19), S(260, 637)) is consistent with cleavage of these sites by cathepsin which is known to cleave S(10). The enhanced cleavage of these sites following addition of camostat in Vero E6 cells suggests some increased diversion of S and N down a cathepsin-dependent cleavage pathway under conditions which inhibit TMPRSS2 and trypsin. While this cell line lacks TMPRSS2 expression, clearly there is some alteration of proteolytic activity following camostat treatment, suggesting inhibition of other proteases in this system which may have a compensatory or complementary function.

The same data lacking normalisation to total viral protein levels can be seen in Fig. S5H. As expected given the lack of significant changes to total N and S protein levels, all sites highlighted in the previous paragraph maintained their direction of change relative to mock, and remained t-test significant in this unnormalised dataset (p < 0.05).

As a majority of these cleavage sites within viral proteins are novel, we lack a functional understanding of their role in viral infection. We sought to examine the importance of several of the novel cleavage sites found within the spike glycoprotein by mutating residues proximal to the cleavage sites and assessing their functions in a pseudovirus entry assay, utilising pseudotyped lentiviral vectors. Given our observed cleavage at Q14 we generated a S13A mutation, altering the P1 residue in this cleavage site following the nomenclature of Schechter and Berger (35). For the cleavage sites at 637 and 671, given that cathepsin is known to cleave spike (10), and indeed the site at 637 showed sensitivity to calpeptin, a calpain and cathepsin inhibitor, we sought to modify cleavage through mutagenesis of the P2 residue within this cleavage site as the P2 site is considered important for cathepsin cleavage (36). This approach generated V635G and C671G mutants. RatG13 spike, a mutant lacking the FURIN cleavage site due to a deletion of four residues (Δ681-684, ΔPPRA) was included as an additional control (37).

In HEK-ACE2 cells, both the V635G and RatG13 mutants showed significantly increased cell entry compared to wild-type (t-test p<0.01), while the C671G mutant showed significantly decreased cell entry (Unpaired Welch’s t-test, p<0.0001)Fig. 3C. Entry for both the S13A and the double V635G/C671G mutant was not significantly different to wild-type Fig. 3C. Representative FACS plots and the gating strategy applied can be found in Fig. S7.

The same pattern was observed in HEK-ACE2-TMPRSS2 cells, though the enhanced cell entry seen for the the V635G and RatG13 mutants did not reach significance Fig. 3D. This may reflect reduced importance of the V635G mutation in cells bearing high levels of TMPRSS2. Similarly while the pattern of partial recovery of the double V635G/C671G mutant remained visible, its entry remained significantly reduced (Unpaired Welch’s t-test, p < 0.05) compared to wildtype Fig. 3D. While reproducible across multiple independent viral stocks and experiments, the phenotypes are modest (2-fold change) in both cell types, except for the marked infectivity defect of the C671G mutant. Representative FACS plots and the gating strategy applied can be found in Fig. S8.

Western blotting of purified pseudovirus particles confirmed expression and incorporation of spike Fig. 3E. Notably con-structs containing the C671G mutation showed limited incorporation and defective spike processing. While this could be in part due to cleavage, this cysteine residue is identified in a disulphide bond in several crystal structures suggesting that a more likely explanation for the lower entry phenotype in pseudovirus bearing this mutation is down to defective protein folding or stability (38), Fig. S9A. The wild-type, S13A and V635G mutants all show S0 (uncleaved) and S1 (cleaved) spike Fig. 3E. Notably the V635G mutant showed significantly increased levels of the cleaved S1, with a near 3:1 ratio of S1 to S0 compared to the wild-type where this is 1:1 Fig. 3F (Unpaired Welch’s t-test, p < 0.05). This could reflect either enhanced cleavage at this location, or increased incorporation of the cleaved V635G S1 into pseudovirus particles. Of note, the increased incorporation of cleaved spike in V635G mutant particles was consistent with the increased infectivity of this mutant Fig. 3C, Fig. S9B.

### SARS-CoV-2 infection induces proteolytic cleavage of multiple host proteins

Examination of cellular neo-N-termini from infected cells, has the potential to indicate activation of host cell proteases, or the targeting of specific host pathways. Motif analysis from both cell lines revealed strong enrichment of serine residues on the P’ region of cleavage sites corresponding to neo-N-termini Fig. 4, AB. Diminished abundance of R/K was also observed, though it should be noted that we excluded neo-N-termini from our analysis which were preceded by R/K and could potentially be arti-facts generated by tryptic digestion during sample processing.

**Fig. 4.**
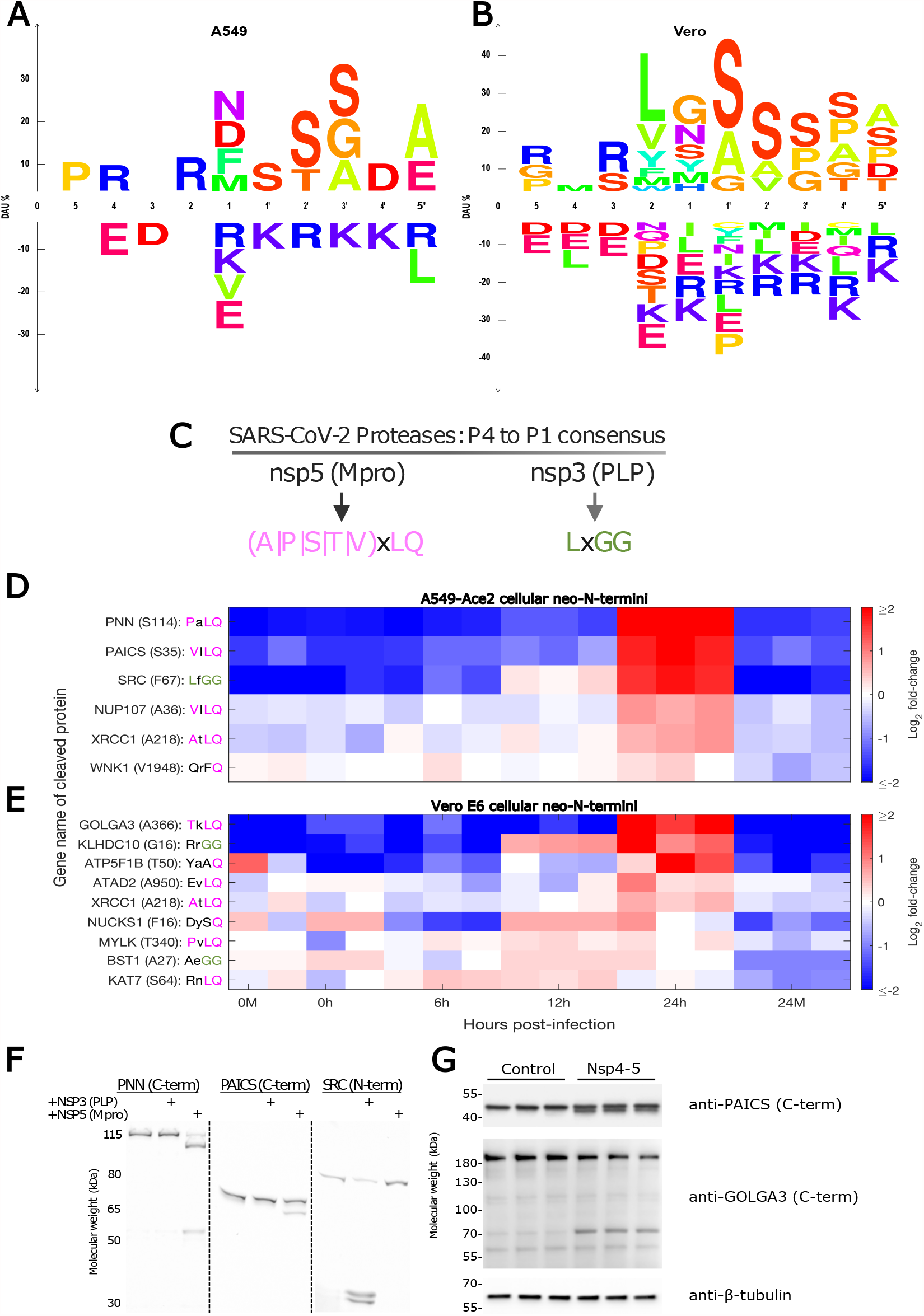
Increased abundance of novel cellular neo-N-termini consistent with SARS-CoV-2 protease consensus sequences suggests viral protease activity on cellular substrates. a) and b) show motif analysis highlighting enriched amino acids proximal to the N-terminus of neo-N-termini enriched in SARS-CoV-2-infected A549-Ace2 and Vero E6 cells respectively. DAU = Differential amino acid usage. The P5 to P5’ positions for the cleavage sites are shown following the nomenclature of Schechter and Berger. c) Consensus motifs for Mpro and PLP. d) and e) show the relative abundance of cellular neo-N-termini identified as significantly upregulated (n = 3 biological replicates, unpaired two sample t-test, multiple-hypothesis corrected q <= 0.05) and matching or resembling the Mpro or PLP consensus motifs from A549-Ace2 or Vero E6 cells respectively. Sequence match to the consensus is indicated by the pink or green coloring of the P4 to P1 positions of the relevant cleavage sites indicating match to the Mpro or PLP P4, P2 or P1 positions respectively. f) *In vitro* validation of GFP-tagged PNN, PAICS and SRC cleavage by SARS-CoV-2 Mpro and PLP, following incubation with 10µM of the respective protease. g) Cell-based validation of Mpro cleavage of GOLGA3 and PAICS following transfection of SARS-CoV-2 Nsp4-5 plasmid. β-tubulin is included as a loading control.

Given this strong enrichment for specific residues in our datasets, we sought to identify if enriched or diminished neo-N-termini could be associated with a causal protease using TopFIND 4.0, which links neo-N-termini to causal proteases (39). As this software does not currently support African Green Monkey datasets, this analysis could be performed only for the Ace2-A549 dataset, the results of which are included in Fig. S10. Of those significantly-regulated neo-N-termini, causal proteases were identified for a subset, though the analysis did not find the overall activities of these causal proteases to be significantly regulated in infected cells at 24h post-infection compared to mock-infected cells (p <= 0.05, Adjusted Fisher’s exact test) S8, though this may reflect low numbers of annotated substrates for individual proteases.

Finally, we performed gene ontology analysis of proteins from which the significantly-regulated neo-N-termini were derived, identifying significantly regulated biological functions, cellular compartments and molecular functions within each cell line dataset Fig. S11, Fig. S12. While many of the enriched GO terms were unique to each cell line, common themes included mitochondria, and the cytoskeleton/cell adhesion.

We then sought to identify neo-N-termini that could be targeted directly by the SARS-CoV-2 proteases, rather than host cell proteases. The consensus sequences for coronavirus proteases are conserved between coronaviruses, with PLP recognising a P4 to P1 LxGG motif, and Mpro recognising a (A|P|S|T|V)xLQ motif (40) Fig. 4C). No strong preference has been identified for either protease at the P3 residue (Fig. 4A). Analysis of both datasets showed strong enrichment for neo-N-termini consistent with cleavage at Mpro motifs (two-tailed Kolmogorov-Smirnov test, p<0.001, Fig. S13A,B). However, no comparable enrichment could be seen for neo-N-termini consistent with cleavage at PLP motifs (Fig. S13C,D). This may reflect fewer cellular protein substrates of PLP compared to Mpro, or higher background levels of neo-N-termini generated by cellular proteases with similar P4 to P1 cleavage specificities as PLP.

Neo-N-termini matching, or close to the consensus sequences, for either Mpro or PLP and showing significant upregulation (t-test, q *≤* 0.05 after correction for multiple hypothesis testing) at 24h post-infection compared to the 24h mock sample were selected for further analysis. Perfect matches to the consensus sequences from A549-Ace2 cells included NUP107, PAICS, PNN, SRC and XRCC1. GOLGA3 and MYLK (MCLK) were identified from Vero E6 cells. Hits from both cell lines that resembled, but did not completely match the consensus sequence were ATAD2, ATP5F1B, BST1, KAT7, KLHDC10, NUCKS1 and WNK1 (Fig. 4D, E). Adding confidence to these observations, approximately half of these hits were also identified in a recent SARS-CoV-2 proximity labelling study (ATP5F1B, GOLGA3, NUP107, PNN, SRC and WNK)(7), and GOLGA3 was additionally identified in an interactome study as an nsp13 interaction partner (6).

SRC, MYLK and WNK are all protein kinases, one of the protein families best studied as drug targets (41). MYLK is especially interesting as dysregulation of MYLK has been linked to acute respiratory distress syndrome -one of the symptoms of severe COVID-19 disease (42). NUP107 is a member of the nuclear pore complex, with nucleocytoplasmic transport a frequent target for viral disregulation (43). GOLGA3 is thought to play a role in localisation of the Golgi and Golgi-nuclear interactions, and was identified in two recent studies of SARS-CoV-2 interactions (6, 7). PNN is a transcriptional activator, forming part of the exon junction complex, with roles in splicing and nonsense-mediated decay. The coronavirus mouse hepatitis virus has previously been shown to target nonsense mediated decay, with pro-viral effects of inhibition (44). PAICS and BST1 both encode enzymes with roles in ADP ribose and purine metabolism respectively, with PAICS previously identified as binding the influenza virus nucleoprotein (45).

The majority of these neo-N-termini showed enrichment at 24h, with levels remaining largely unchanged at earlier timepoints, especially for Mpro substrates (Fig. 4D,E). This matches the timing for peak viral RNA, protein expression and titres over the timepoints examined (Fig. 1B-D). Exceptions to this trend include the potential PLP substrates, 2/3 of which begin to show increased abundance at 12h post-infection, with BST1 appearing to peak at 12h rather than 24h, indicating a potential temporal regulation of the two viral proteases. Data for all quantified and filtered N- and neo-N-termini from A549-Ace2 and Vero E6 cells is available in tables S4 and S5 respectively. Paired analysis of enriched neo-N-termini together with our unenriched dataset, allowed inference of cleavage stoichiometry for a subset of these significantly enriched cleaved cellular neo-N-termini by the HIquant approach (46) Fig. S14, suggesting that by 24h in SARS-CoV-2-infected cells, the majority of these proteins are present in the cleaved form.

We sought to validate a subset of prospective SARS-CoV-2 protease substrates *in vitro*, using the L. tarentolae system previously used to identify cleavage of proteins involved in the immune response by the SARS-CoV-2 proteases (14). For these assays, the target protein is fused (N-or C-terminally) to GFP, which is then imaged directly in the SDS-PAGE gel. This system sucessfully validated cleavage of PNN, PAICS and SRC by Mpro (PNN, PAICS) and PLP (SRC) respectively (Fig. 4F). It also indicated additional cleavage products of PNN, and SRC not identified in the original mass spectrometry study. SRC cleavage was identified by N-terminomics following a LfGG motif yielding a neo-N-terminus at F67 (Fig. 4F). Two SRC cleavage products migrate at slightly over 30kDa (including the GFP tag). The second cleavage product found in vitro migrates at slightly higher molecular weight, consistent with cleavage at an LaGG motif 19 amino acids downstream of the first cleavage site, generating a neo-N-terminus at V86 (Fig. 4,H).

Titration of the amount of PLP included in the reaction resulted in dose-dependent cleavage of SRC (Fig. S15).

In addition to a cleavage product consistent with the cleavage at S114 observed in the N-terminomics migrating between the 80-115kDa markers, a second cleavage product of PNN was also identified, migrating at slightly over 50kDa including the GFP tag (Fig. 4H). However there are multiple candidate cleavage sites located in this portion of PNN that could explain this cleavage event.

We also validated cleavage of PAICS and GOLGA3 by SARS-CoV-2 in a cell-based assay. To generate functional Mpro, a plasmid containing the nsp4-5 sequence was generated, permitting autocleavage at the nsp4-5 junction and generation of an authentic N-terminus for nsp5 (Mpro). Transfection of HEK 293 cells with this construct resulted in cleavage of PAICS and GOLGA3. Both antibodies recognise the C-terminus of the cleaved proteins. As with the *in vitro* assay, cleavage of PAICS resulted in the appearence of a single cleavage product. This assay recognises endogenous rather than tagged PAICS which is why both uncleaved and cleaved PAICS have differing apparent molecular weights in Fig. 4F and G. Anti-GOLGA3 cleavage results in the appearence of a single-cleavage product at slightly over 70kDa. This may reflect further proteolytic cleavage of this protein given that the observed cleavage at 365/366 would be expacted to result in a 40k reduction in the apparent molecular weight of GOLGA3 which migrates at slightly over its predicted molecular weight of 167kDa.

### Prospective MPro and PLP substrates are necessary for efficient viral replication, and represent targets for pharmacological intervention

To investigate if the putative cellular substrates of MPro and PLP identified in the N-terminomic analyses are necessary for efficient viral replication, an siRNA screen was conducted Fig. 5. Where proteolytic cleavage inactivates cellular proteins or pathways inhibitory for SARS-CoV-2 replication, siRNA depletion would be anticipated to result in inreased viral titres and/or RNA levels. If proteolysis results in altered function that is beneficial for the virus, we would expect siRNA depletion to result in a reduction in viral titres/RNA levels. Proteins with neo-N-termini showing statistically significant increased abundance during SARS-CoV-2 infection and either matching, or similar to the viral protease consensus sequences were selected for siRNA depletion.

**Fig. 5.**
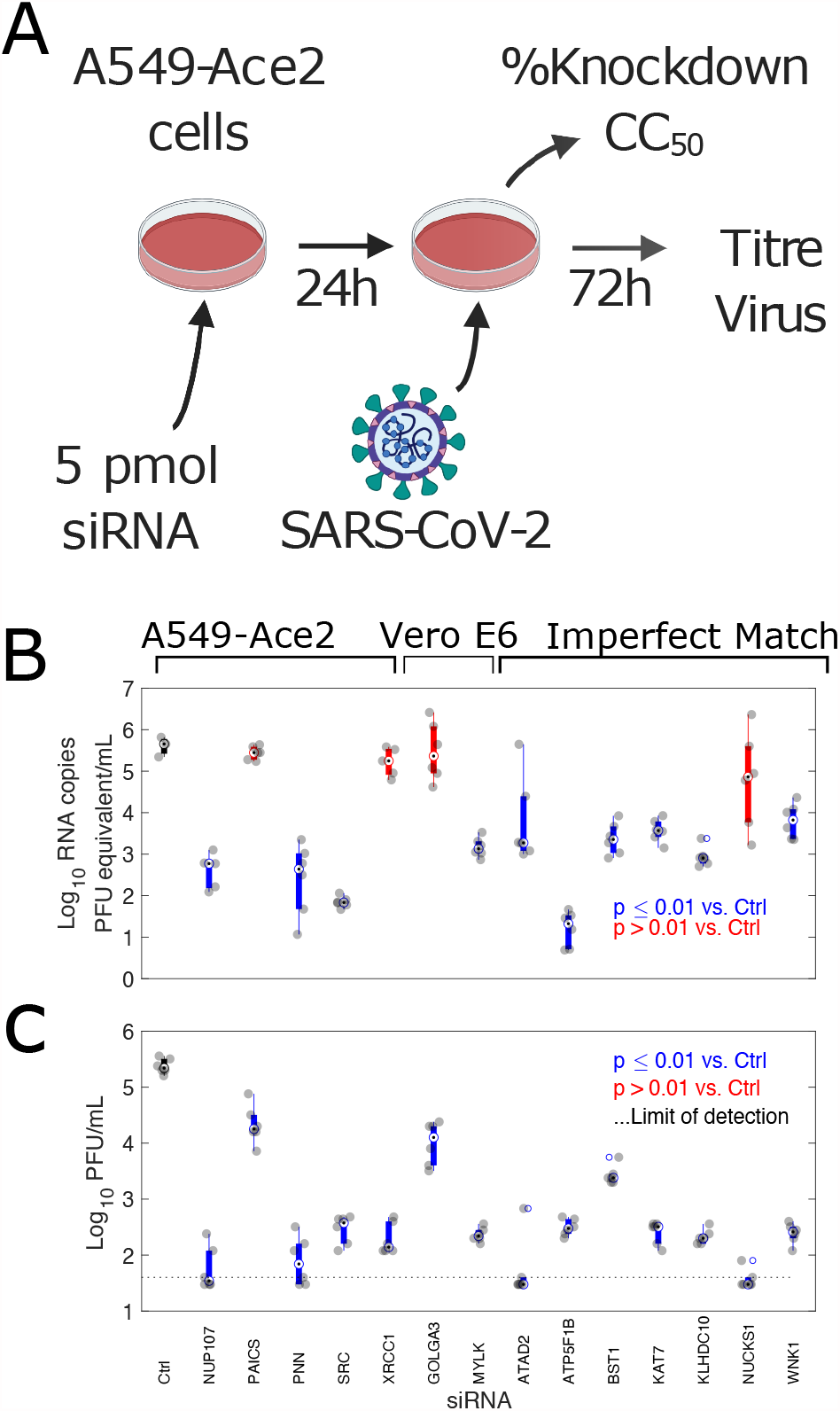
siRNA depletion of potential MPro and PLP substrates results in significant reductions to viral RNA copies and titres in A549-Ace2 cells. A) Experimental design, B) Viral RNA copies and C) titres in supernatant at 72h post-infection, following infection 24h post-transfection with the indicated siRNA (n *≥* 4 biological replicates). Colored bars indicate median (hollow circle with dot) and 25th and 75th percentiles. Individual datapoints are shown in grey. Significance was calculated by one-way ANOVA. Blue bars indicate samples with significantly reduced viral RNA copies or titres (p<=0.01). Those not meeting this threshold are shown in red. The control siRNA-treated sample is indicated with a black bar. The limit of detection in the plaque assay was calculated to be 40 PFU/mL (dotted line).

Infection of A549-Ace2 cells was performed 24h post-transfection with the indicated siRNA and allowed to proceed for 72h (Fig. 5A). Cell viability for all targets was comparable to untreated controls (Fig. S16). siRNA knock-down efficiency at the time of infection was confirmed by qRT-PCR (Fig. S17), with a low of 77% efficiency for NUCKS1, and averaging over 95% efficiency for most targets. 10/14 coronavirus protease substrates showed significant reductions (one-way ANOVA, p *≤* 0.01) in viral RNA levels, averaging a 100-1000-fold median decrease in viral RNA equating to pfu equivalents per ml at 72h post-infection compared to treatment with a control siRNA (Fig. 5B). PAICS, GOLGA3, NUCKS1, and XRCC1 did not show asignificant drop in RNA copy number following siRNA treatment. Plaque assays were then conducted on these samples to determine whether this observed reduction in viral RNA levels reflected a reduction in infectious virus titres (Fig. 5C). All 14 potential substrates showed a statistically significant (one-way ANOVA, p *≤* 0.01) reduction in viral titres following siRNA depletion. For PAICS and GOLGA3, which did not show reduced RNA levels, these reductions were approximately 10-fold. Most other siRNA targets showed reduced titres in the 100-1000-fold range. These differences in outcome between viral RNA levels and plaque assays may result from a subset of proteins required for efficient viral replication. While efficient mRNA knockdown was shown for all targets Fig. S17, it is also possible that this discrepancy between viral RNA levels and titres may result from differences in protein half-life of the knockdown targets. This could result in proteins with longer half lives only giving a phenotype at later stages of infection when infectious virus is produced. Finally, we extended our analysis to two previously-identified SARS-CoV-2 substrates (14), IRF3 and TAB1 Fig. S18 not identified in our N-terminomic analysis. We found that in line with previous reports (47), the antiviral IRF3 resulted in an aproximately 3-fold increase in viral titres and increased RNA copies. In contrast, TAB1 depletion resulted in reduced SARS-CoV-2 RNA copies and viral titres, possibly reflecting the role of TAB1 in negative regulation of antiviral responses (48). In line with the apparently pro-viral nature of the majority of the prospective SARS-CoV-2 substrates, we observed in a microscopy-based viability assay that siRNA-treated infected cells appeared to show enhanced viability compared to infected cells treated with a scrambled siRNA control, though only reached statistically significance (One-way ANOVA, Tukey’s correction for multiple hypothesis testing) for some substrates (PNN, XRCC1, GOLGA3, ATP5F1B).

A subset of the prospective viral protease substrates have commercially-available inhibitors, notably SRC and MYLK. In the case of SRC these include tyrosine kinase inhibitors in current clinical use. In light of the siRNA screening results we concluded that pharmacological inhibition of SARS-CoV-2 protease substrates could represent a viable means to inhibit SARS-CoV-2 infection. Dose-response experiments were conducted with 7 inhibitors to determine whether pharmacological inhibition of SARS-CoV-2 protease substrates could be employed as a potential therapeutic strategy (Fig. 6; Fig. S19). Of these, two tyrosine kinase inhibitors: Bafetinib and Sorafenib showed inhibition at concentrations which did not result in cytotoxicity in the human cell line A549-Ace2 (Fig. 6). In the case of Bafetinib, a Lyn/Bcr-Abl inhibitor which has off-target activity against SRC the *IC*50 was in the nanomolar range (*IC*50: 0.79 µM, 95% confidence interval 0.23-1.35 µM). Bafetinib has recently been independently identified as an inhibitor of the coronaviruses OC43 and SARS-COV-2 in a large-scale drug-repurposing screen (49). Inhibition with Sorafenib which was included as a positive control and does not directly target any of the protease substrates was in the low micromolar range (Fig. 6), in line with a previously published report (4). Two inhibitors were trialed against MYLK. These were MLCK inhibitor peptide 18, and ML-7. Only ML-7 showed inhibition of SARS-CoV2, with inhibition in the low micromolar range (*IC*50: 1.7 µM, 95% confidence interval 1.51-1.80 µM), at concentrations which did not induce cytotoxicity (Fig. 6; Fig. S20). ML-7 and MLCK inhibitor peptide 18 have different mechanisms of action, with MLCK inhibitor peptide 18 outcompeting kinase substrate peptides, and ML-7 inhibiting ATPase activity. All four had *CC*50 values over the 10µM maximum concentration tested, except ML-7 which had a *CC*50 of 5 µM. Bafetinib did show reduced viability at the two highest concentrations tested (10 µM, 3.3 µM), though not reaching 50% reduction (Fig. S20).

**Fig. 6.**
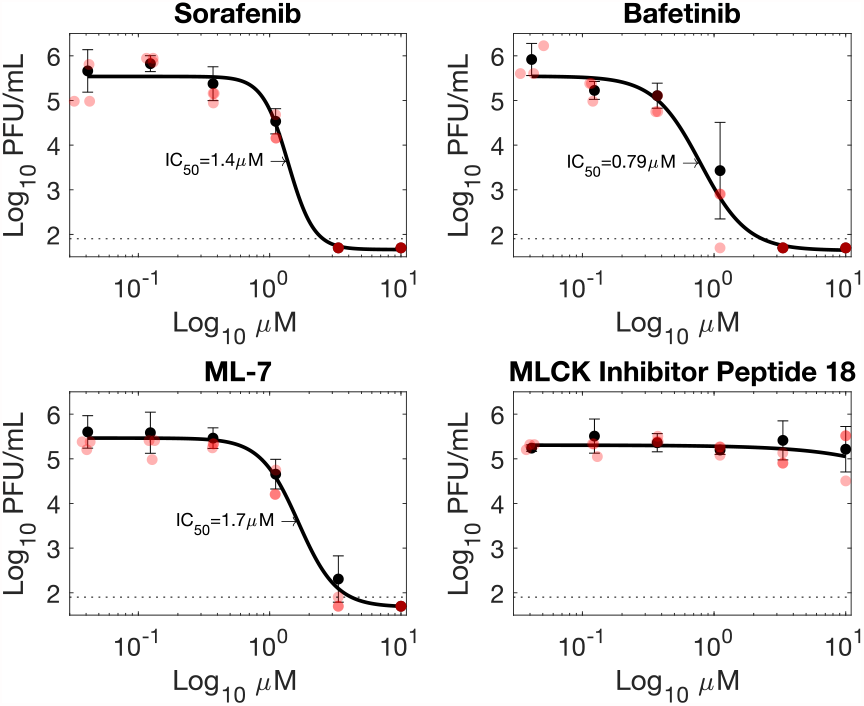
Inhibitors targeting viral protease substrates reduce SARS-CoV-2 titres in A549-Ace2 cells. Sorafenib is a tyrosine kinase inhibitor previously shown to inhibit SARS-CoV-2 replication (4). Bafetinib is a dual ABL/LYN inhibitor with off-target activity against SRC kinase. ML-7 and MLCK target myosin light chain kinase (MYLK/MLCK). Black circles and error bars represent mean and standard deviation from 3 biological replicates. Red circles indicate individual datapoints. PFU = plaque-forming units.

The other 3 tyrosine kinase inhibitors tested (Bosutinib, Saracatinib, Dasatinib) all showed inhibition Fig. S19, however cytotoxicity results obtained with the assay used were also high preventing the unambiguous determination of whether inhibition was specific or due to cytotoxicity Fig. S20. However, it should be noted that these agents have been reported to be cytostatic in A549 cells, and the CellTiter Gloassay used to assess viability measures cellular metabolism so will not distinguish between cytostatic and cytotoxic effects.

## Discussion

We employed a mass spectrometry approach to study proteolytic cleavage events during SARS-CoV-2 infection. Substrates of viral proteases are frequently inferred through studies of related proteases (50). However, such approaches are unable to identify novel substrates, and even closely-related proteases can differ in their substrate specificity (51). Mass spectrometry-based approaches to identify protease substrates by identifying the neo-N-terminal peptides generated by protease activity have existed for a number of years (22, 52–54), however, they have seen only limited application to the study of viral substrates (55, 56), and have not been previously applied to the study of proteolysis during coronavirus infection.

While our approach identified multiple novel viral and cellular cleavage sites, it also failed to identify multiple known cleavage sites, including the FURIN cleavage site in spike, and multiple cleavage sites within the viral polyprotein. This can be understood from the dependence of the approach on the specific protease used for mass spectrometry analysis. Isobaric labelling prior to trypsin digestion blocks tryptic cleavage at lysine residues and causes trypsin to cleave solely after arginine residues. This results in the generation of long peptides and if the specific cleavage site does not produce a peptide of suitable length for analysis (typically 8-30 amino acids) then it will be missed. This can be alleviated through the application of multiple mass spectrometry-compatible proteases in parallel, yielding multiple peptides of different length for each cleavage site (57, 58). This would both increase the number of sites identified and cross-validate previously identified cleavage sites. These methods will likely prove a fruitful avenue for future investigations of proteolysis during infection with SARS-CoV-2 and other viruses that employ protease-driven mechanisms of viral replication.

Our approach identified multiple cleavage sites within viral proteins. In some cases, such as the nucleocapsid protein, cleavage by cellular proteases has been observed for SARSCoV (18, 19), though the number of cleavage products observed was much higher in our study (Fig. 2). Compared to the gel-based approaches used in the past, our approach is much more sensitive for detecting when protease activity results in N-termini with ragged ends, due to further exoproteolytic activity. Examples of this in our data are particularly evident in Fig. 2 for the nucleocapsid and ORF3a where neo-N-termini appear in clusters. Cleavage sites within the nucleocapsid and spike protein are of particular interest as these are the two viral antigens to which research is closely focuses for both testing and vaccination purposes. In this context, neo-N-termini are of interest as N-termini can be recognised by the immune response, as they are typically surface-exposed. Antibodies recognising neo-N-termini such sites will not be detected in tests using complete or recombinant fragments that do not account for such cleavage sites. Indeed, a recent study revealed altered antigenicity of proteolytic proteoforms of the SARS-CoV-2 nucleocapsid following autolysis (26), and loss of antibody responses to spike following mutations in circulating SARS-CoV-2 variants altering signal peptide cleavage (34). Understanding cleavage events can also inform interpretation of protein structural analysis, for example in the ORF3a viroporin (27). Knowledge of cleavage sites can permit further analysis of spike entry mechanisms, and vaccine design, and inform genotype-to-phenotype predictions.

Formation of the most prominent neo-N-terminus we identified in SARS-CoV-2 spike at 637 appeared dependent on cathepsins and/or calpains, as its appearance was limited by calpeptin treatment. Mutation of the P2 residue in the putative cleavage site, resulting in mutant V365G, led to an increased incorporation of cleaved spike in pseudotyped viral particles. Consistent with an increased content of fusion-competent cleaved spike, the V635G mutant showed an increased infectivity in HEK-Ace2 target cells.

Why blocking the formation of the 637 neo-N-terminus promotes cleaved spike incorporation remains to be elucidated. A possible explanation may lie in a competition between different cleavage sites in the producer cell, with cleavage at 637 inhibiting cleavage at the FURIN site, or inhibiting the incorporation of spike trimers already cleaved at the FURIN site. The capacity of producer cells to cleave viral glycoproteins at alternative sites may thus be viewed as an intrinsic defense mechanism. In contrast, the capacity of viral proteases to cleave multiple host proteins, as demonstrated in this study, contributes to well-established mechanisms aiming at inhibiting innate host responses, including in particular the interferon pathway (14). Therefore, the diversity of proteolytic cleavage events revealed by N-terminomics may reflect another layer in the dynamic evolutionary conflict between viruses and their hosts.

Proteolytic cleavage can alter protein function in several ways, including inactivation, re-localisation, or altered function including the removal of inhibitory domains. Our siRNA screen showed knockdown of the majority of potential protease targets we identified was inhibitory to SARS-CoV-2 replication (Fig. 5). Indeed, with the exception of the previously reported antiviral protein IRF3, no siRNA treatment resulted in higher viral titres or RNA levels, suggesting that inactivation is not the prime purpose of these cleavage events. This suggests that in many cases, proteolytic cleavage by viral proteases may be extremely targeted, serving to fine-tune protein activity, rather than merely serving as a blunt instrument to shut down unfavorable host responses. It is also worth noting the low overlap between our infection-based study, and a subsequently released N-terminomics dataset which used incubation of cell culture lysates with recombinant Mpro (59). Only GOLGA3 was common to both studies, in spite of the larger number of cleavage events identified in the Koudelka *et al*. study. However in such a lysate-based experiment subcellular compartmentalisation and regulation of relative enzyme and substrate localisation & abundance is lost, so can risk identifying cleavage events not possible *in vivo* during genuine infection. An improved understanding of the exact ways in which proteolytic cleavage is regulated, modulates protein activity, and serves to benefit viral replication will be crucial for targeting cellular substrates of viral proteases as a therapeutic strategy.

## Limitations

In this study, we used two cell line models to characterise the effects of SARS-CoV-2 infection on protease activity and the generation of viral and cellular cleavage products. Notably, we tested the efficiency of several inhibitors against SARS-CoV-2 infection only in the context of the A549-Ace2 cell line model. Given previous experiences translating cell culture findings for SARS-CoV-2 to the clinic, it is important to note these results present preliminary data that must be further validated in other models, *in vivo*, and through clinical trials before use in patients for the treatment of COVID-19 disease.

## Materials & Methods

### Cell culture & Virus

Vero E6 (Vero 76, clone E6, Vero E6, ATCC® CRL-1586TM) authenticated by ATCC and tested negative for mycoplasma contamination prior to commencement were maintained in Dulbecco’s modified Eagle’s medium (DMEM; Thermo Fisher Scientific) containing 10% (v/v) fetal bovine serum (FBS, ThermoFisher Scientific) and penicillin/streptavidin (ThermoFisher Scientific). A549-Ace2 cells, a human lung epithelial cell line that over-expresses ACE2, were kindly provided by Oliver Schwartz (Institut Pasteur) (60). A549-Ace2 cells were cultured in DMEM supplemented with 10% FBS, penicillin/streptavidin and 10 *µ*g/ml blasticidin (Sigma) and maintained at 37°C with 5% CO2. The SARS-CoV-2 isolate BetaCoV/France/IDF0372/2020 was supplied through the European Virus Archive goes Global (EVAg) platform. Viral stocks were prepared by propagation in Vero E6 cells in DMEM supplemented with 2% FBS. For protease inhibitor experiments employing calpeptin and camostat mesylate, these drugs or an equal volume of vehicle (DMSO) were supplemented to the medium 12h post-infection at 50mM final concentration. All experiments involving live SARS-CoV-2 were performed in compliance with Institut Pasteur Paris’s guidelines for Biosafety Level 3 (BSL-3) containment procedures in approved laboratories. All experiments were performed in at least three biologically independent samples. For spike-pseudotyped lentivector production and infections, HEK293Tn were maintained in Dulbecco’s modified Eagle medium (DMEM) supplemented with 10% fetal bovine serum and 100µg/mL penicillin/streptomycin (complete medium), and cultured at 37°C under 5% CO2. HEK293T-hACE2-TMPRSS2 (called HEK-ACE2-TMPRSS2) with inducible TPMRSS2 expression were a gift from Julian Buchrieser and Olivier Schwartz (60). These cells were maintained in complete medium with blasticidin (10 µg/mL, InvivoGen) and puromycin (1 µg/mL, Alfa Aesar), and TM-PRSS2 expression was induced by addition of doxycycline (0.5 µg/mL, Sigma).

### SARS-CoV-2 titration by plaque assay

Vero E6 cells were seeded in 24-well plates at a concentration of 7.5×104 cells/well. The following day, serial dilutions were performed in serum-free MEM media. After 1 hour absorption at 37°C, 2x overlay media was added to the inoculum to give a final concentration of 2% (v/v) FBS / MEM media and 0.4% (w/v) SeaPrep Agarose (Lonza) to achieve a semi-solid overlay. Plaque assays were incubated at 37° C for 3 days. Samples were fixed using 4% Formalin (Sigma Aldrich) and plaques were visualized using crystal Violet solution (Sigma Aldrich).

### Infections for N-terminomic/proteomic analysis

N-terminomic sample preparation is based around Weng et al. 2019 Mol. Cell. Proteomics, adapted for TMTpro-based quantitation (22, 61). A protocol for this TMTpro-adapted method can be found at https://www.protocols.io/view/tmtpro-hunter-n-terminomics-bi44kgyw. Vero E6 or A549-Ace2 cells were seeded using 2×106 cells in T25 flasks. The following day cells were either mock infected or infected with SARS-CoV-2 at a MOI of 1 in serum-free DMEM at 37°C for 1 hour. After absorption, the 0 hour samples were lysed immediately, while the media for other samples was replaced with 2% FBS / DMEM (ThermoFisher Scientific) and incubated at 37°C for times indicated before lysis. Cells were washed 3x with PBS (ThermoFisher Scientific) before lysing them in 100 mM HEPES pH 7.4 (ThermoFisher Scientific), 1% Igepal (Sigma Aldrich), 1% sodium dodecyl sulfate (SDS; ThermoFisher Scientific), and protease inhibitor (mini-cOmplete, Roche). Samples were then heated to 95°C for 5 minutes, before immediately freezing at -80°C. Samples were then thawed and incubated with benzonase for 30 min at 37°C. Sample concentrations were normalized by BCA assay, and 25µg of material from each sample was used for downstream processing.

DTT was added to 10mM and incubated at 37°C for 30 min, before alkylation with 50mM 2-chloroacetamide at room temperature in the dark for 30 min. DTT at 50mM final concentration was added to quench the 2-chloroacetamide for 20 min at room temperature. Samples were washed by SP3-based precipitation (62). Each sample was resuspended in 22.5µL 6M GuCl, 30µL of 0.5M HEPES pH8, and 4.5µL TCEP (10mM final) and incubated for 30 minutes at room temperature.

0.5 mg of individual TMTpro aliquots (Lot VB294905) were resuspended in 62uL of anhydrous DMSO. 57µL of the TMT-pro was then added to each sample, mixed and incubated for 1.5h. Label allocation was randomized using the Matlab Randperm function. Excess TMTpro was quenched with the addition of 13µL of 1M ethanolamide and incubated for 45 min. All samples were combined for downstream processing. SP3 cleanup was performed on the combined samples. These were resuspended in 400µL of 200mM HEPES pH8, containing Trypsin gold at a concentration of 25ng/µL and incubated overnight at 37°C.

Samples were placed on a magnetic rack for 5 min. 10% of the samples was retained for the unenriched analysis. The remaining material was supplemented with 100% ethanol to a final concentration of 40%, undecanal added at an undecanal:peptide ratio of 20:1 and sodium cyanoborohydride to 30mM. pH was confirmed to be between pH7-8 and the samples were incubated at 37°C for 1h. Samples were then sonicated for 15 seconds, and bound to a magnetic stand for 1 min. The supernatant was retained and then acidified with 5% TFA in 40% ethanol. Macrospin columns (Nest group) were equilibrated in 0.1% TFA in 40% ethanol. The acidified sample was applied to the column, and the flow through retained as the N-terminal-enriched sample.

Both unenriched and enriched samples were desalted on macrospin columns (Nest group), before drying down again. Off-line basic reverse phase fractionation for both unenriched and enriched samples was performed on a Waters nanoAcquity with an Acquity UPLC M-Class CSH C18 130A 1.7µm, 300µm x 150µm column. The sample was run on a 70 minute gradient at 6µL/min flow rate. Gradient parameters were 10 min 3% B, 10-40 min 3-34% B, 40-45 min 34-45% B, 45-50 min 45-99%B, 50-60 min 99% B, 60.1-70 min 3% B. Buffers A and B were 10mM ammonium formate pH10, and 10mM ammonium formate pH10 in 90% acetonitrile respectively. Both samples were resuspended in buffer A, and 1 minute fractions were collected for 1-65 min of the run. These were concatenated into 12 (1:13:24…) or 5 fractions (1:6:11…) for unenriched and enriched samples respectively using a SunChrom Micro Fraction Collector. Samples were dried down and resuspended in 1% formic acid for LC-MS/MS analysis.

### Mass spectrometry

LC-MS/MS analysis was conducted on a Dionex 3000 coupled in-line to a Q-Exactive-HF mass spectrometer. Digests were loaded onto a trap column (Acclaim PepMap 100, 2 cm x 75 microM inner diameter, C18, 3 microM, 100 Å) at 5 µL per min in 0.1%(v/v) TFA and 2%(v/v) acetonitrile. After 3 min, the trap column was set inline with an analytical column (Easy-Spray PepMap® RSLC 15 cm x 50cm inner diameter, C18, 2 microlM, 100 Å) (Dionex). Peptides were loaded in 0.1%(v/v) formic acid and eluted with a linear gradient of 3.8–50% buffer B (HPLC grade acetonitrile 80%(v/v) with 0.1%(v/v) formic acid) over 95 min at 300 nl per min, followed by a washing step (5 min at 99% solvent B) and an equilibration step (25 min at 3.8% solvent). All peptide separations were carried out using an Ultimate 3000 nano system (Dionex/Thermo Fisher Scientific).. The Q-Exactive-HF was operated in data-dependent mode with survey scans aquired at a resolution of 60,000 at 200m/z over a scan range of 350-2000m/z. The top 16 most abundant ions with charge states +2 to +5 from the survey scan were selected for MS2 analysis at 60,000 m/z resolution with an isolation window of 0.7m/z, with a (N)CE of 30%. The maximum injection times were 100ms and 90ms for MS1 and MS2 respectively, and AGC targets were 3e6 and 1e5 respectively. Dynamic exclusion (20 seconds) was enabled.

### Data analysis

All data were analysed using Maxquant version 1.6.7.0 (63). Custom modifications were generated to permit analysis of TMTpro 16plex-labelled samples. FASTA files corresponding to the reviewed Human proteome (20,350 entries, downloaded 8th May 2020), and African Green monkey proteome (Chlorocebus sabeus, 19,223 entries, downloaded 16th May 2020). A custom fasta file for SARS-CoV-2 was generated from the Uniprot-reviewed SARS-CoV-2 protein sequences (2697049). This file was modified to additionally include the processed products of pp1a and pp1b, novel coding products identified by ribo-seq (25), as well as incorporate two coding changes identified during sequencing (spike: V367F, ORF3a: G251V). All FASTA files, TMT randomisation strategy, and the modifications.xml file containing TMTpro modifications have been included with the mass spectrometry data depositions. Annotated spectra covering peptide N-termini of interest were prepared using xiS-PEC v2(64).

Several different sets of search parameters were used for analysis of different experiments.

### For analysis of unenriched material from fractionated lysates

Default MaxQuant settings were used with the following alterations. As the experimental design meant unenriched samples contained a majority of peptides lacking N-terminal TMT labelling, quantification was performed at MS2-level with the correction factors from Lot VB294905 on lysine labelling only with the N-Terminal label left unused. Digestion was trypsin/p with a maximum of 3 missed cleavages. Carbamidomethylation of cysteines was selected as a fixed modification. Oxidation (M), Acetylation (Protein N-terminus), and N-terminal TMTpro labelling were selected as variable modifications. PSM and Protein FDR were set at 0.01.

### For analysis of fractionated, N-terminally-enriched material

Default MaxQuant settings were used with the following alterations. Quantification was performed at MS2-level with the correction factors from Lot VB294905. Digestion was semi-specific ArgC, as TMTpro labelling of lysines blocks trypsin-cleavage. Carbamidomethylation of cysteines was selected as a fixed modification. Oxidation (M), Acetylation (Protein N-terminus), Gln/Glu to pyroglutamate were selected as variable modifications. PSM and Protein FDR were set at 0.01.

### For analysis of viral protein neo-N-termini from fractionated, N-terminally-enriched material

Default MaxQuant settings were used with the following alterations. MS1-based quantitation was selected. Digestion was ArgC, sei-specific N-terminus. Carbamidomethylation of cysteines was selected as a fixed modification. Oxidation (M), Acetylation (Protein N-terminus), Gln/Glu to pyroglutamate, and TMTpro modification of N-termini and lysine residues were selected as variable modifications. PSM and Protein FDR were set at 0.01.

All downstream analysis was conducted in Matlab, external packages used include BreakYAxis (65). Reverse hits and contaminants were removed, peptides were filtered to meet PEP *≤* 0.02. For quantitative analysis, peptides were further filtered at PIF *≥*0.7. TMTpro data was normalised for differences in protein loading by dividing by the label median, rows were filtered to remove rows with more than 2/3 missing data. Missing data was KNN imputed, and individual peptides were normalised by dividing by their mean abundance accross all TMTpro channels. As the objective was to identify protein cleavage events, peptides were further filtered to remove those beginning at the first or second amino acid in a protein sequence that represent the native N-terminus. +/-methionine. neo-N-termini were annotated if they matched known signal peptides. For non-quantitative analysis (e.g. mapping of viral neo-N-termini), peptides were filtered to retain only blocked (acetylated, TMTpro labelled, and pyroglutamate) N-termini. pyroglutamate-blocked N-termini were discarded if they were preceeded by arginine or lysine as these could represent artifactual cyclization of tryptic N-termini. Fractional protein or peptide intensity was calculated as the total intensity for the protein or peptide, multiplied by the fraction of the summed normalised TMTpro intensity represented by a particular TMTpro label of interest.

Inference of cleaved and uncleaved proteoforms of protease substrates was performed using the HIquant approach (46), using the unenriched and N-terminally enriched data, processed as above. To ensure that unenriched tryptic peptides represented a simple, rather than complex mixture of proteoforms, inference was performed for cellular substrates where the unenriched tryptic peptides formed a single cluster based on the Euclidean distance using the Matlab evalcluster (‘linkage’) implementation.

Visualisation of the Y636/S637 cleavage site within the SARS-CoV-2 spike glycoprotein structure was performed using PDB: 6×6P (28), in UCSF ChimeraX v1.0 (66).

### Data analysis for motif analysis, association of causal proteases and GO enrichment

Data was normalised for loading by dividing each TMT intensity column by the column median. The rows with 75% or more missing values were removed. The remaining missing values were imputed using K-nearest neighbour algorithm. Each row was then normalised by the row mean. Peptides that required imputation of more than one value per group were excluded. Neo-N-peptides were selected by the following criteria: start position at amino acid > 2, not among known signal peptides, not among predicted signal peptides, not preceded by arginine or lysine. Prediction of potential signal peptides was performed using SignalP v5.0 (67) tool -only peptides with prediction confidence greater than 0.9 and predicted cleavage site located 5 or less amino acids from a cleavage site of a detected peptide were considered.

All analysis described in this section was performed using the R statistical programming environment (68). Differential abundance analysis was performed using limma (69) package. The statistical significance of the results was assessed using Storey’s q-values (24) (q-value <= 0.05). Gene set enrichment analysis was performed using clusterProfiler (70) package in R using BioSigDB (71, 72) gene sets. Significant pathways selected by p-values, adjusted for multiple testing using Benjamini-Hochberg method (p-value <=0.05). Peptide motif analysis was done using the dagLogo (73) package. Peptide regions of 5 amino acids before and after the cleavage site were selected and motif analysis was performed using Fisher’s exact test (p-value <=0.05). All figures were produced using package ggplot2 (74). Data analysis was performed by members of the University of Liverpool Computational Biology Facility.

### Production of spike-pseudotyped lentivectors

Lentiviral particles encoding the SARS-CoV-2 spike were prepared by transient transfection of HEK293Tn cells using the CaCl_2_ method. The lentiviral vector pCDH-EF1a-GFP (System Bioscience), the packaging plasmid psPAXII (Addgene), the spike expression vector phCMV-SARS-CoV-2-Spike (a gift from O. Schwartz), and the pRev plasmid (a gift from P. Charneau) were mixed at a 2:2:1:1 ratio and transfected at 252 µg DNA per 175 cm2 cell flask. The pQCXIP-Empty plasmid was used as a negative control for spike expression. At 48h after transfection, supernatants were collected and concentrated by ultracentrifugation at 23,000 g for 1h 30m at 4°C on a 20% sucrose cushion. Viral particles were resuspended in PBS and frozen in aliquots at -80°C until use. Gag p24 antigen concentration was measured with the Alliance HIV-1 p24 Antigen ELISA kit (Perkin Elmer).

Point mutations to generate the mutants were introduced by site-directed mutagenesis of phCMV-SARS-CoV-2-Spike using Q5 polymerase (Thermo Scientific) and validated by sanger sequencing. The primers used were: S13A F: GTG TCC GCT CAG TGC GTG AAC CTG ACC ACA C, S13A R: CAC TGA GCG GAC ACC AGT GGC AGC AGC ACC, V635G F: CGC GGG TAC TCC ACC GGC AGC AAT GTG, V635G R: GTA CCC GCG CCA TGT TGG TGT CAA TTG ATC, C671G F: ATC GGC GCC TCC TAT CAG ACC CAG ACC, C671G R: GGC GCC GAT TCC GGC TCC GAT GGG GAT ATC.

### Infection with spike-pseudotyped lentivectors

The day before infection, 100,000 HEK-ACE2 or HEK-ACE2-TMPRSS2 cells were plated in 96-well plates and TMPRSS2 was induced by the addition of doxycycline. HEK-ACE2 +/-TMPRSS2 were infected with the equivalent of 2 µg of p24 Gag for each spike lentivector, in final volume of 100 µL. Infection was quantified by measuring the percentage of GFP+ cells two days post-infection by flow cytometry. Cells were harvested, washed in PBS, and stained with the viability dye eF780 (eBioscience) for 30 min at 4°C. After two washes in PBS, cells were fixed with paraformaldehyde 2% (ThermoFisher) and acquired on an Attune NxT flow cytometer. Results were analyzed with FlowJo software (v10.7.1), with statistical analyses carried out with the GraphPad Prism software (v9).

### Western blotting of spike-pseudotyped lentivectors

To prepare protein extracts, cells were lysed in buffer with NaCl 150 mM, Tris HCl 50 mM (pH8), 1% Triton, EDTA 5 mM, supplemented with protease inhibitors (Roche) for 30 min on ice. For lentiviral particle extracts, an equivalent of 500 ng of p24 Gag was lysed in buffer with 1% Triton (ELISA kit, Alliance Perkin Elmer) for 30 min on ice. To preserve antibody reactivity, samples were not heated nor reduced before being run in a 4-12% acrylamide denaturing gel (NP0323, NuPAGE, ThermoFisher), and then transferred onto a nitrocellulose membrane (IB23001, ThermoFisher). The membrane was blocked with 5% dried milk in PBS Tween 0.1%, before incubation with the primary antibody for 1h at RT, followed by 3 washes, and incubation with the secondary antibody for 30 min at RT. After 3 more washes, the fluorescent signal was revealed on a LiCor Odyssey 9120 imaging system. Images were quantified with the ImageStudioLite (v5.2.5) software, using a mode with automated background subtraction. Primary antibodies consisted in the human anti-spike mAb 48 (1:1,000; a gift from H. Mouquet) or the mouse anti-p24 Gag MAB7360 (R&D Systems; 1:1000). Anti-human or mouse IgG secondary antibodies, conjugated to DyLight-800 (A80-304D8, Bethyl Laboratories) or DyLight-680 (SA5-35521, ThermoFisher) respectively, were used at a 1:10,000 concentration.

### In vitro cleavage assays

*In vitro* cleavage assays were performed using the Leishmania tarentolae (LTE) system as described (14). SRC, PAICS, PNN and RPA2 (control) were cloned as GFP fusion proteins into dedicated Gateway vectors for cell-free expression. Open Reading Frames (ORFs) in pDonor were sourced from the Human ORFeome collection, version 8.1 and transferred into Gateway destination vectors that include N-terminal (SRC) or C-terminal (PAICS, PNN and RPA2) Fluorescent proteins. The specific Gateway vectors were created by the laboratory of Pr. Alexandrov and sourced from Addgene (Addgene plasmid # 67137; http://n2t.net/addgene:67137; RRID:Addgene_67137). LTE extracts for *in vitro* expression were prepared in-house as described (75). Purified recombinant Mpro and PLP were generated by the UNSW protein production facility as described previously (14).

The SRC, PAICS, PNN and RPA2proteins were expressed individually in 10 µL reactions (1µL DNA plasmid at concentrations ranging from 400ng/L to 2000ng/L added to 9 µL of LTE reagent). The mixture was incubated for 30 minutes at 27°C to allow the efficient conversion of DNA into RNA. The samples were then split into controls and protease-containing reactions. The proteases PLpro (nsp3) and 3CLpro (nsp5) were added at various concentrations, and the reactions were allowed to proceed for another 2.5h at 27°C before analysis. The controls and protease-treated LTE reactions were then mixed with LDS (Bolt LDS Sample Buffer, ThermoFisher) and loaded onto SDS-page gels (4-12% Bis-Tris Plus gels, ThermoFisher); the proteins were detected by scanning the gel for green (GFP) fluorescence using a ChemiDoc MP system (BioRad) and proteolytic cleavage was assessed from the changes in banding patterns. Note that in this protocol, the proteins are not treated at high temperature with the LDS and not fully denatured, to avoid destruction of the GFP fluorescence. As proteins would retain some folding, the apparent migration on the SDS-page gels may differ slightly from the expected migration calculated from their molecular weight. We have calibrated our SDS-page gels and ladders using a range of proteins, as shown previously (14).

### Transfection and cell-based validation of proteolytic cleavage by Western blotting

A mammalian expression plasmid expressing the coding sequence of SARS-Cov-2 Nsp4-Nsp5 in a pCDNA3.1 backbone was synthesized (GeneArt™ Gene synthesis, ThermoFisher, USA). HEK 293T cells in a 6 well plate were transfected with polyethylenimine (PEI) and 2ug of Nsp4-Nsp5 fusion construct, or with pCDNA3.1 control, in 3 biological replicates. After 48 hours, cells were lysed with RIPA buffer (ThermoFisher, USA) in presence of Phosphatase/Protease inhibitors cocktail (ThermoFisher, USA). The lysate was centrifuged (15 min at 4°C and 13000rpm) and the supernatant was collected. Protein concentration was quantified using Pierce™ BCA Protein Assay kit (ThermoFisher, USA) and Western Blot performed with standard protocol. Briefly, proteins were seperated on a precast 4-20% gradient gel (Biorad, USA) and transferred on a nitrocellulose membrane using a semi-dry Trans-Blot Turbo Transfer System and Trans-Blot Turbo Transfer Buffer (Biorad, USA). Membranes were blocked for 1 hour with 5% milk in TBST (Tris-Buffered Saline and Tween 20) buffer, rinsed, and incubated overnight at 4°C with primary antibodies in 2% BSA in TBST. Membranes were washed with TBST and incubated for 2h at room temperature with Horseradish peroxidase (HRP)-linked secondary secondary antibody (Cell signaling #7074). Chemiluminescent signal was revealed using SuperSignal™ West Pico PLUS Substrate (ThermoFisher, USA) and imaged with an Azure 600 Imaging system (Azure Biosystem, USA). The primary antibodies used were PAICS (Bethyl A304-547A-T), GOLGA3 (Bethyl A303-404A-T) and β-Tubulin (Cell Signaling #2128). Primary and secondary antibodies were used at 1/1000 and 1/5000 dilutions, respectively. Importantly, primary antibodies against PAICS and GOLGA3 recognized C-terminal immunogens, ensuring that cleaved proteins could be detected.

### Virus infections in siRNA-based cellular protein knockdowns

Host proteins were knocked-down in A549-Ace2 cells using specific dsiRNAs from IDT. Briefly, A549-Ace2 cells seeded at 1×104 cells/well in 96-well plates. After 24 hours, each well was transfected with 5 pmol of individual dsiRNAs using Lipofectamine RNAiMAX (Thermo Fisher Scientific) according to the manufacturer’s instructions. 24 hours post transfection, the cell culture supernatant was removed and replaced with virus inoculum (MOI of 0.1 PFU/cell). Following a 1 hour adsorption at 37°C, the virus inoculum was removed and replaced with fresh 2% FBS/DMEM media. Cells were incubated at 37°C for 3 days before supernatants were harvested. Samples were either heat-inactivated at 80°C for 20 min and viral RNA was quantified by RT-qPCR, using previously published SARS-CoV-2 specific primers targeting the N gene (76). RT-qPCR was performed using the Luna Universal One-Step RT-qPCR Kit (NEB) in an Applied Biosystems QuantStudio 7 thermocycler, using the following cycling conditions: 55 °C for 10 min, 95 °C for 1 min, and 40 cycles of 95 °C for 10 sec, followed by 60 °C for 1 min. The quantity of viral genomes is expressed as PFU equivalents, and was calculated by performing a standard curve with RNA derived from a viral stock with a known viral titer. Alternatively, infectious virus titers were quantified using plaque assays as described above. To quantify siRNA-based cellular protein knockdowns, A549-Ace2 cells were seeded and transfected with individual dsiRNAs as described above. After 24 hours incubation at 37 °C cells were lysed and RNA was extracted using Trizol (ThermoFisher Scientific) followed by purification using the Direct-zol-96 RNA extraction kit (Zymo) following the manufacturer’s instructions. RNA levels of target proteins were subsequently quantified by using RT-with the Luna Universal One-Step RT-qPCR Kit (NEB) in an Applied Biosystems QuantStudio 7 thermocycler using gene-specific primers. Expression levels were compared to scram-bled dsiRNA-transfected cells und normalized to expression of human beta-actin. Knockdown efficiencies were calculated using ΔΔCt in Matlab.

To assess cell viability after siRNA knockdowns, cells were seeded and transfected as described above. 24 hours after transfection cell viability was measured using alamar-Blue reagent (ThermoFisher Scientific), media was removed and replaced with alamarBlue and incubated for 1h at 37°C and fluorescence measured in a Tecan Infinite M200 Pro plate reader. Percentage viability was calculated relative to untreated cells (100% viability) and cells lysed with 20% ethanol (0% viability), included in each plate.

For cell counting to determine cell numbers, cells were fixed in formalin to deactivate virus. The fixed cells were stained with 5µg/ml of Hoechst 33258 (Sigma). The assay plates were imaged on an IX-83 automated inverted microscope (Olympus) using a 10x objective. The DAPI settings (Ex UV 377/50, Em 415–480) were used to image Hoechst 33258. The acquisition setup was configured to image 4 sites per well. The nuclei were identified using the object detection module in the ScanR analysis software.

### Drug Screens and Cytotoxicity analysis

Black with clear bottom 384 well plates were seeded with 2×103 A549-Ace2 cells per well. The following day, individual compounds were added using the Echo 550 acoustic dispenser at concentrations indicated 2 hours prior to infection. DMSO-only (0.5%) and remdesivir (10*µ*M; SelleckChem) controls were added in each plate. After the pre-incubation period, the drug-containing media was removed, and replaced with virus inoculum (MOI of 0.1 PFU/cell). Following a one-hour adsorption at 37°C, the virus inoculum was removed and replaced with 2% FBS/DMEM media containing the individual drugs at the indicated concentrations. Cells were incubated at 37°C for 3 days. Supernatants were harvested and heat-inactivated at 80°C for 20 min. Detection of viral genomes from heat-inactivated was performed by RT-qPCR as described above. Cytotoxicity was determined using the CellTiter-Glo luminescent cell viability assay (Promega). White with clear bottom 384 well plates were seeded with 2×103 A549-Ace2 cells per well. The following day, individual compounds were added using the Echo 550 acoustic dispenser at concentrations indicated. DMSO-only (0.5%) and camptothecin (10 *µ*M; Sigma Aldrich) controls were added in each plate. After 72 h incubation, 20*µ*l/well of Celltiter-Glo reagent was added, incubated for 20 min and the luminescence was recorded using a luminometer (Berthold Technologies) with 0.5 sec integration time. Curve fits and *IC*50/*CC*50 values were obtained in Matlab.

## Supporting information

Table S1 - Viral neo-N-termini from A549-Ace2 cells

Table S2 - Viral neo-N-termini from Vero E6 cells

Table S3 - All viral peptides identified in enriched and unenriched experiments for both Vero E6 and A549-Ace2 experiments

Table S4 - N-terminomics TMT quantification data for A549-Ace2 cells

Table S5 - N-terminomics TMT quantitation data for Vero E6 cells

Table S6 - neo-N-termini within 5aa of VOI/VOC characteristic mutations

Table S7 - neo-N-termini within 10aa of VOI/VOC characteristic mutations

Table S8 - TopFIND analysis of causal proteases

Table S10 - Supplemental Resources Table

## Data availability

All mass spectrometry data, database FASTA files, and the matlab scripts used to generate the data in this manuscript can be found on the ProteomeX-change Consortium (http://proteomecentral.proteomexchange.org) via the PRIDE repository (77), and on GitHub respectively. Specifically the proteomics datasets have been deposited as described in table S9, where reviewer usernames and passwords are provided. The Matlab scripts used to process the mass spectrometry data and produce the figures in this manuscript have been tested in Matlab versions R2019b with the Statistics Machine Learning Toolbox, on Mac OS Catalina. These can be accessed through the Emmott Lab Github page at: https://github.com/emmottlab/sars2nterm/. Other reagent and oligo sequence details are described in table S10.

## ACKNOWLEDGEMENTS

We thank members of the Centre for Proteome Research, especially Rob Beynon and Jos Sarsby, as well as Nikolai Slavov & Aleksandra Petelski (Northeastern University) for constructive comments. We also thank Agnès Zettor and Soizick Lucas-Staat from the Chemogenomic and Biological Screening Platform for their technical assistance. We thank Julian Buchrieser, Olivier Schwartz, Pierre Charneau, Cyril Planchais, and Hugo Mouquet for the gift of reagents. The A549-Ace2, HEK-ACE2 and HEK-ACE2-TMPRSS2 cells were a gift from Olivier Schwartz (Institut Pasteur). The authors would like to thank Katherina Michie, Jack Bennett and key personnel at the protein production facility of UNSW for the purification of PLpro and 3CLpro of SARS-CoV-2. The authors would like to thank Emma Ollivier for the initial in vitro cleavage work on SRC and for useful discussions and comments, and Dominic J.B Hunter for production of cell-free extracts. The authors would also like to thank Prof. Alexandrov for the cell-free plasmids compatible with LTE protein production. This work was supported by the Laboratoire d’Excellence “Integrative Biology of Emerging Infectious Diseases” (grant ANR-10-LABX-62-IBEID) to M.V. S.G. is the recipient of a MESR/Ecole Doctorale BioSPC ED562, Université de Paris fellowship. L.A.C. is supported by Institut Pasteur TASK FORCE SARS COV2 (Tropicoro project), DIM ELICIT Region Ile-de-France, and ANRS. E.E. is supported by startup funding from the University of Liverpool, as well as a Wellcome Trust ISSF Interdisciplinary & Industry Award. E.E. is grateful for the support of GoFundMe donors for sponsoring SARS-CoV-2 research in his laboratory.

## Supplementary Material

**Table S1.** Viral neo-N-termini identified from SARS-CoV-2-infected A549-Ace2 cells - .csv

**Table S2.** Viral neo-N-termini identified from SARS-CoV-2-infected Vero E6 cells - .csv

**Table S3.** All Viral peptides identified accross enriched and unenriched A549-Ace2 and Vero E6 datasets - .csv

**Table S4.** Quantification data for all N- and neo-N-termini quantified from SARS-CoV-2 infected A549-Ace2 cells - .csv

**Table S5.** Quantification data for all N- and neo-N-termini quantified from SARS-CoV-2 infected Vero E6 cells - .csv

**Table S6.** Viral neo-N-termini from A549-Ace2 cells located within 5 amino acids of characteristic mutations from Variants of Interest or Variants of Concern circulating May 2021 - .csv

**Table S7.** Viral neo-N-termini from A549-Ace2 cells located within 10 amino acids of characteristic mutations from Variants of Interest or Variants of Concern circulating May 2021 - .csv

**Table S8.** TopFIND analysis of neo-N-termini significantly increased or decreased in abundance in A549-Ace2 cells (24h Infected/24h Mock) -.xlsx

**Table S9.**
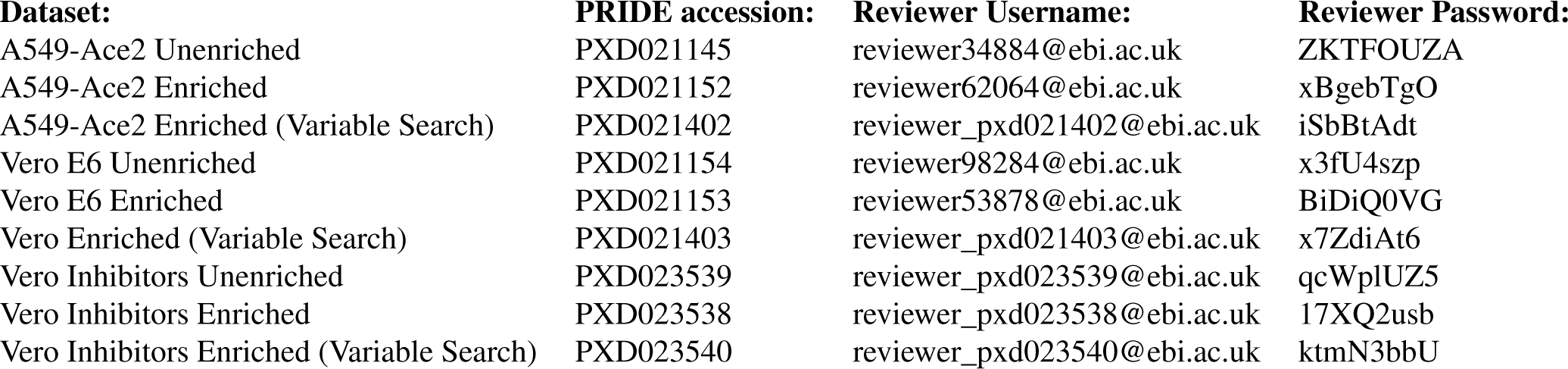
Proteomic datasets and access details

**Table S10.** Oligo sequences and reagent details - .csv

**Fig. S1.**
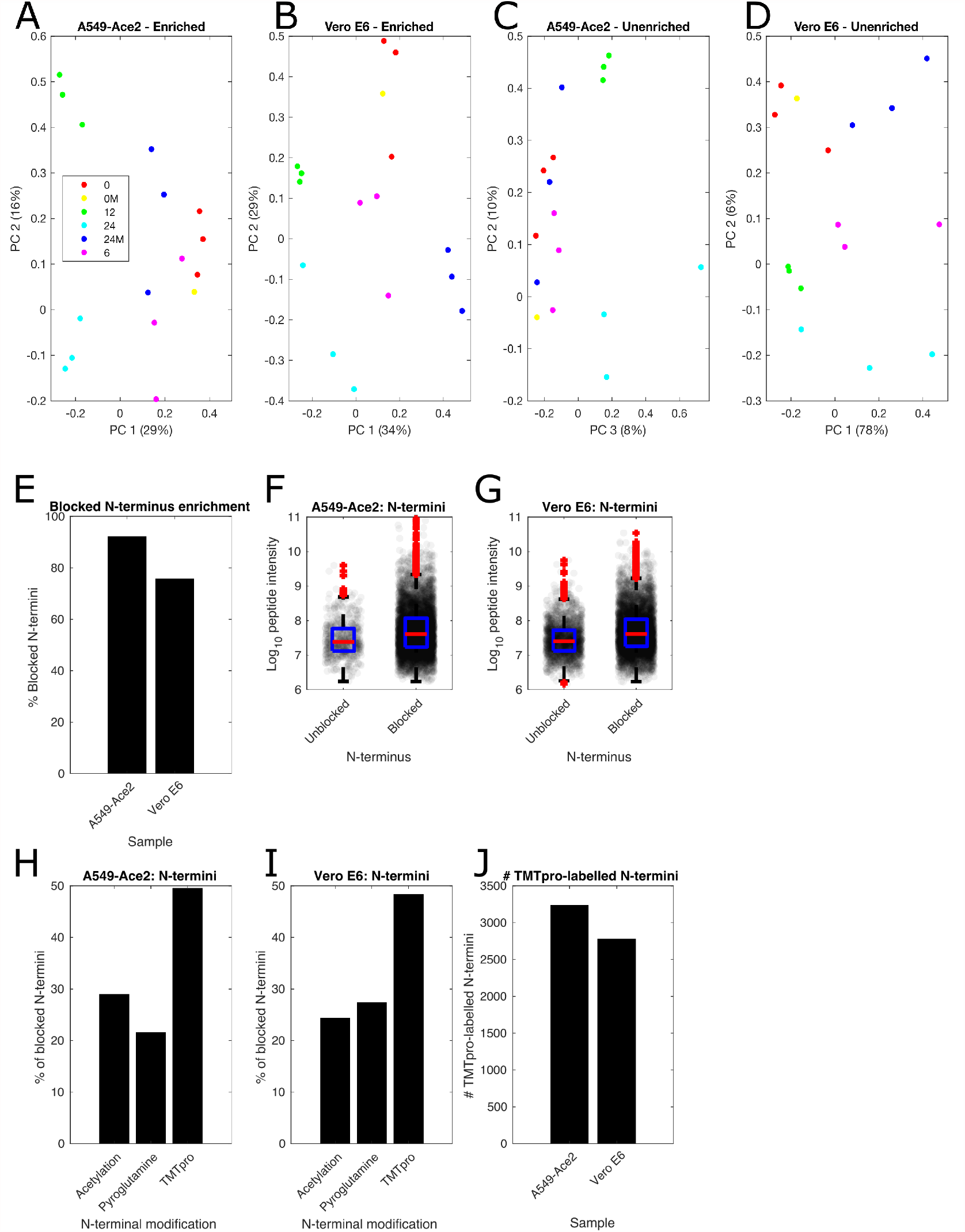
Quality control of the proteomic datasets, both pre- and post-enrichment for N-termini. A-D) Principal component analysis separates infected from mock cells and shows reproducible clustering of biological replicates. E) Enrichment results in a majority of peptide identifications belonging to blocked N-termini, with blocked N-termini most abundant in both F) A549-Ace2 and G) Vero E6 cells. In both cell lines, TMTpro-labelled N-termini are the most abundant enriched N-termini H), I). J) over 2700 TMTpro-labelled N-termini were identified from each dataset.

**Fig. S2.**
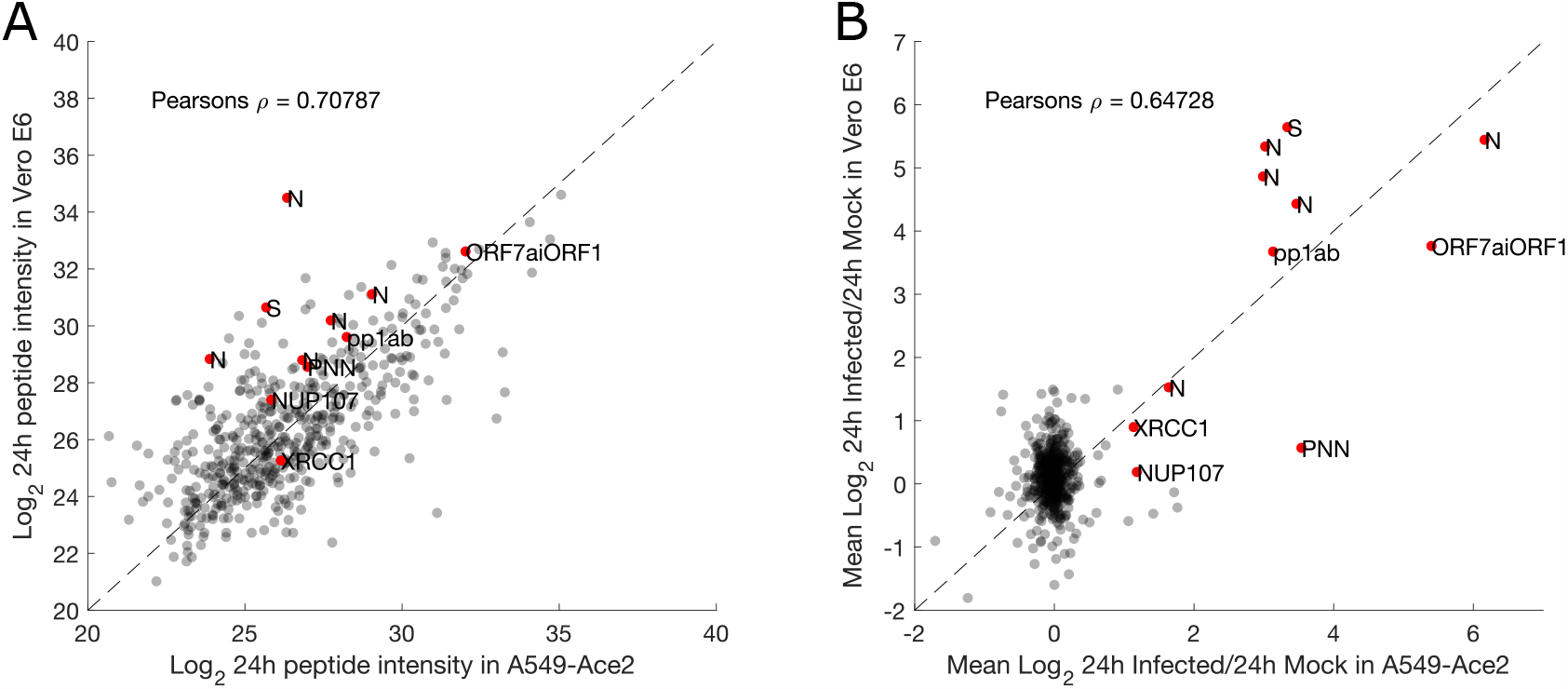
Correlations of neo-N-termini shared between both the A549-Ace2 and Vero E6 datasets. A) Mean peptide intensity correlations for the 24h infected A549-Ace2 and Vero E6 samples. B) *Log*2 24h infected/mock correlations for the A549-Ace2 and Vero E6 samples. Mean values (n=3) are shown, specific neo-N-termini mentioned by name in the text as highlighted in red and labelled. 497 neo-N-termini were quantifiable and common to both datasets.

**Fig. S3.**
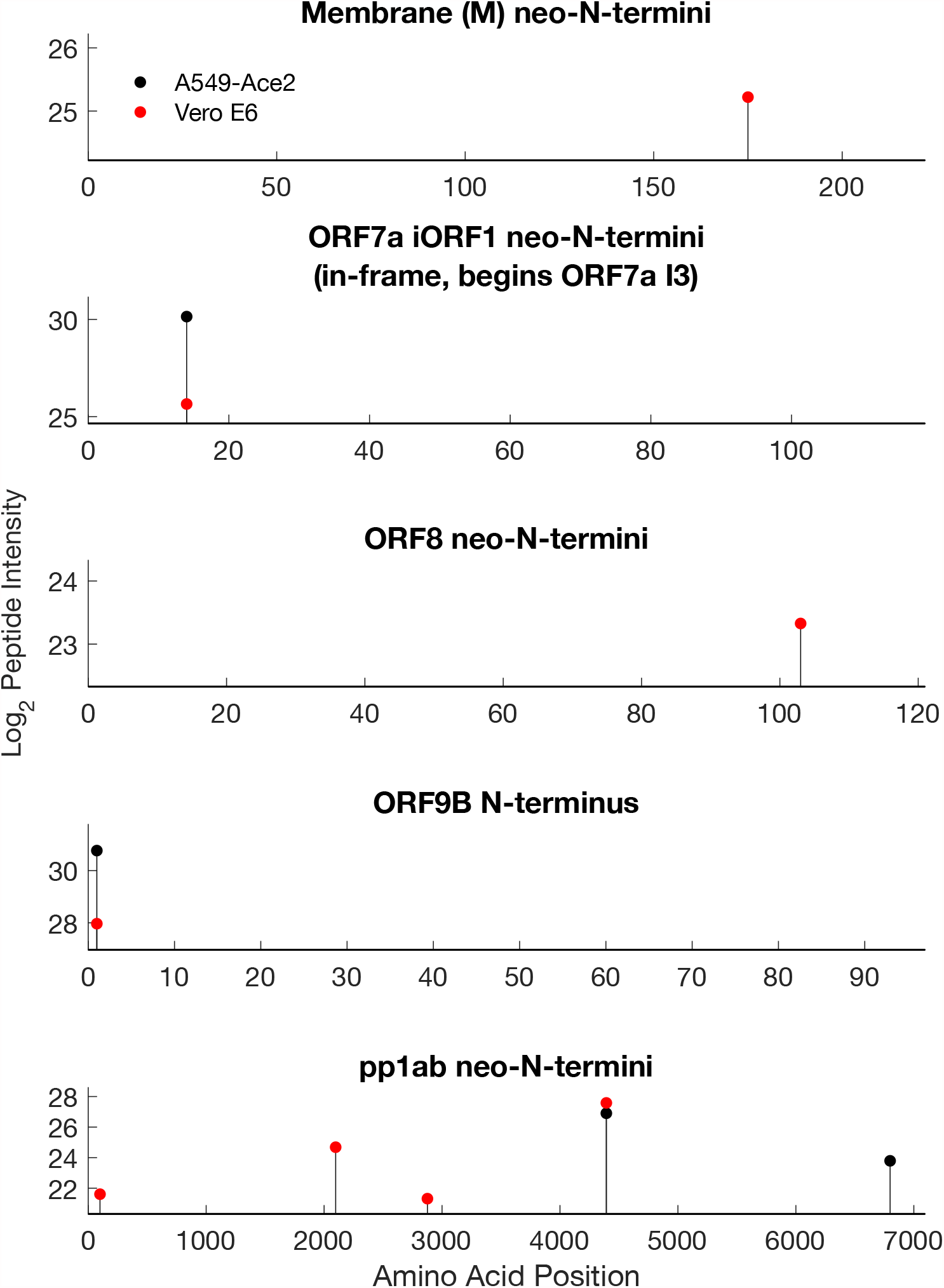
Viral N-termini and neo-N-termini identified from the viral M, ORF7A/ORF7A iORF1, ORF8, ORF9B and pp1ab replicase. Please note that ORF7A iORF1 is an N-terminally truncated form of ORF7A that initiates at isoleucine 3 in the ORF7A sequence. The indicated neo-N-terminus beginning at amino acid 14, would therefore be amino acid 16 in ORF7A.

**Fig. S4.**
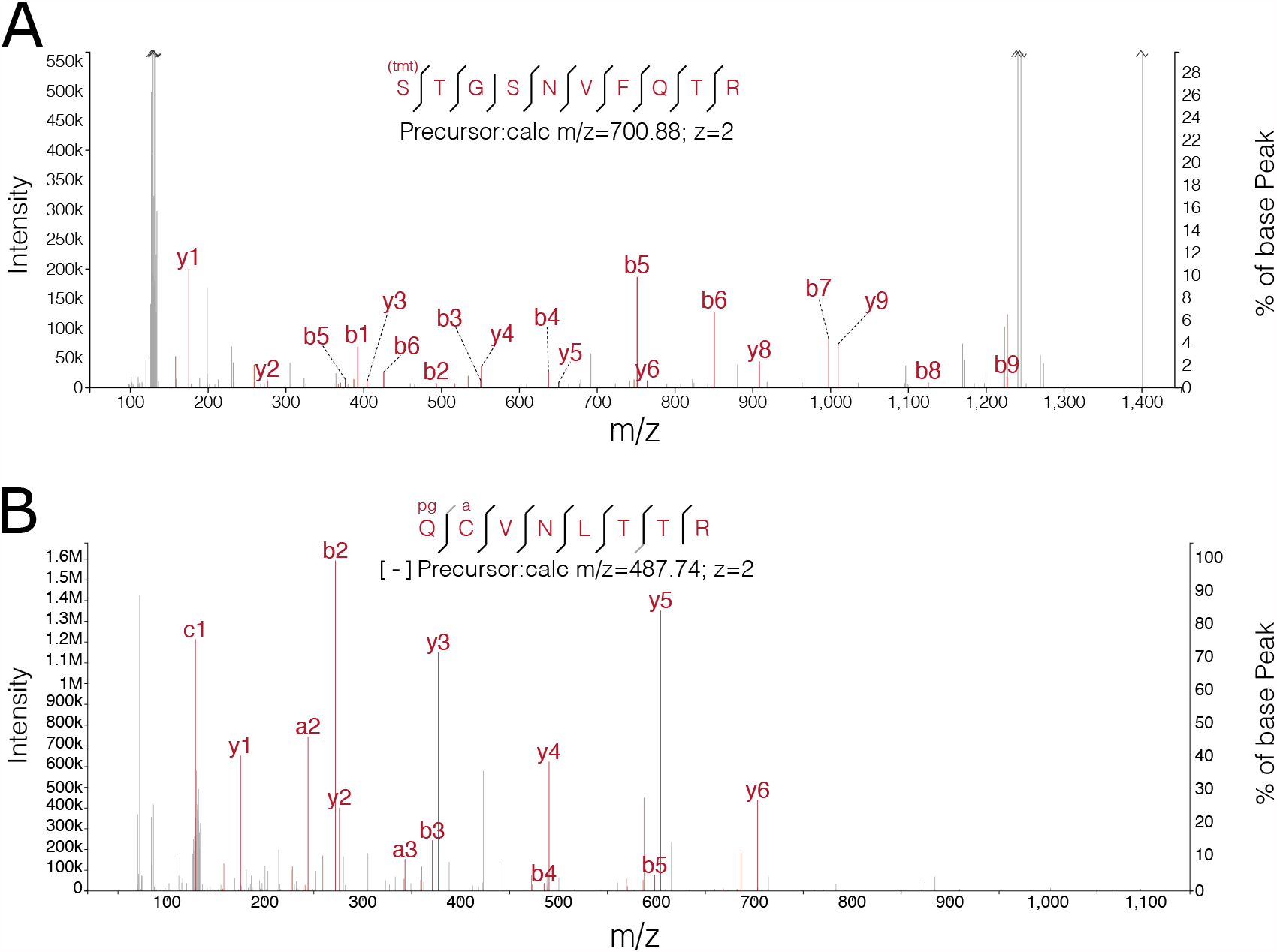
Tandem Mass spectra for a) SARS-CoV-2 spike 637 neo-N-terminus and b) pyroglutamate-modified SARS-CoV-2 spike signal peptide cleavage site beginning Q14. b/y ions are shown in both. a/c ions are also included in b) due to the fragmentation properties of pyroglutamate-containing peptides (78).

**Fig. S5.**
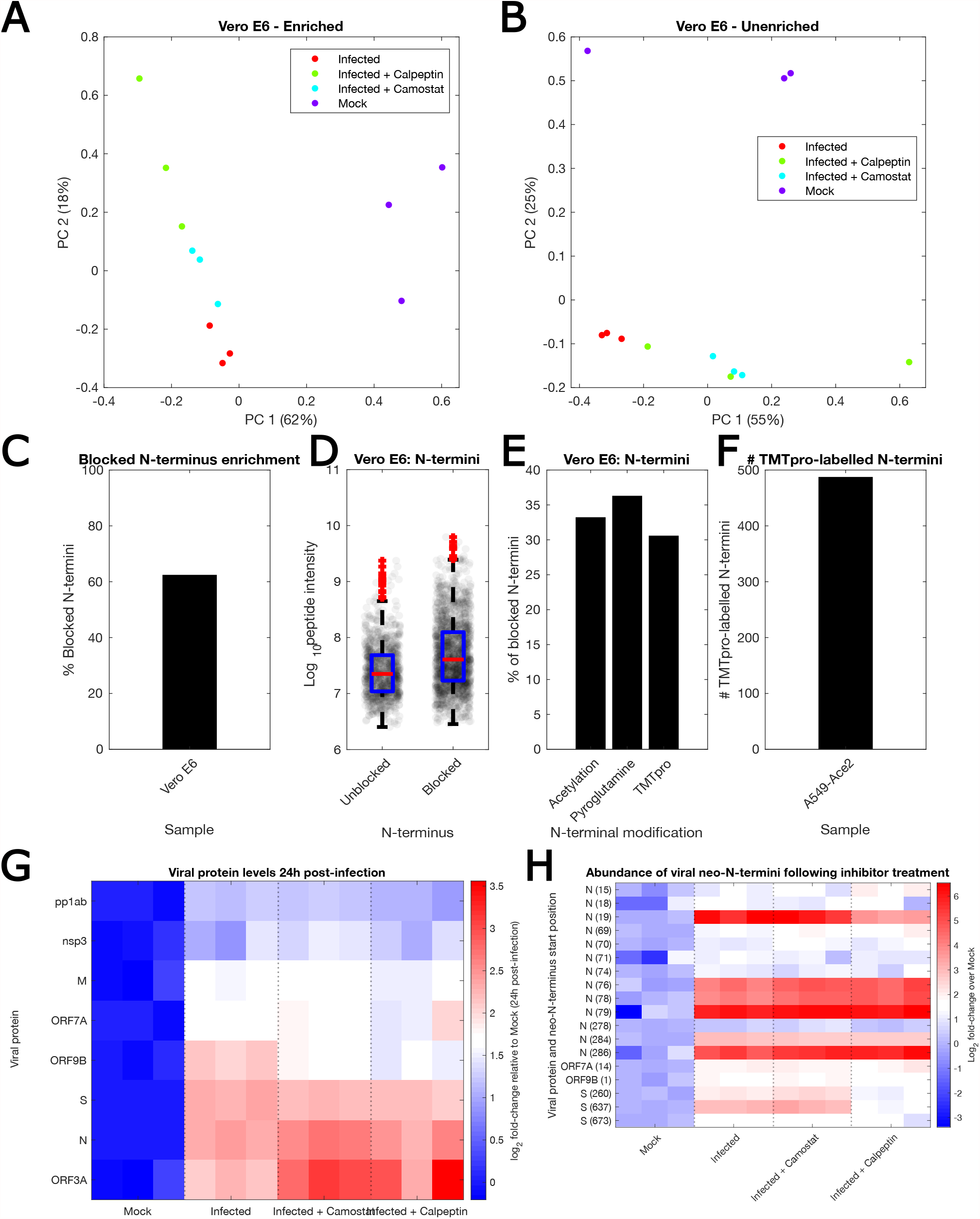
Quality control of the protease inhibitor-treated proteomic dataset, both pre- and post-enrichment for N-termini. A-B) Principal component analysis separates infected from mock cells and shows reproducible clustering of biological replicates. C) Enrichment results in a majority of peptide identifications belonging to blocked N-termini, with blocked N-termini most abundant D). In both cell lines, TMTpro-labelled N-termini are the most abundant enriched N-termini E). F) over 475 TMTpro-labelled N-termini were identified from each dataset. G) Relative abundance of viral proteins in Vero E6 cells mock-or infected with SARS-CoV-2 and infected in the presence of inhibitors (n = 3 biological replicates). H) Relative abundance of neo-N-termini from viral proteins in the same treatment groups. Data is not re-normalised to the total abundance of the viral protein in which the cleavage site is found (n = 3 biological replicates).

**Fig. S6.**
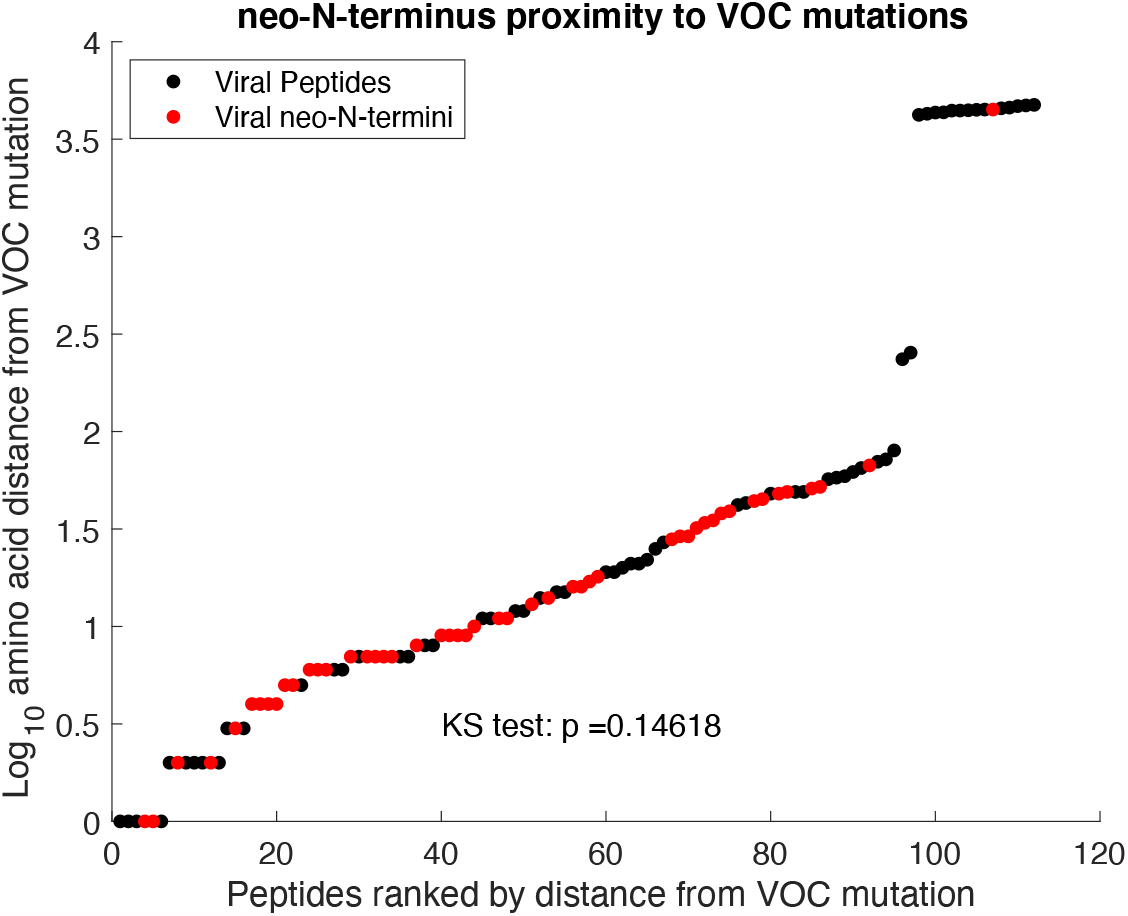
Distance of neo-N-termini (red) from mutations present in Variants Of Concern (VOC) relative to all tryptic peptides identified from viral proteins in unenriched datasets (black). p = ns, one-sample Kolmogorov-Smirnov test.

**Fig. S7.**
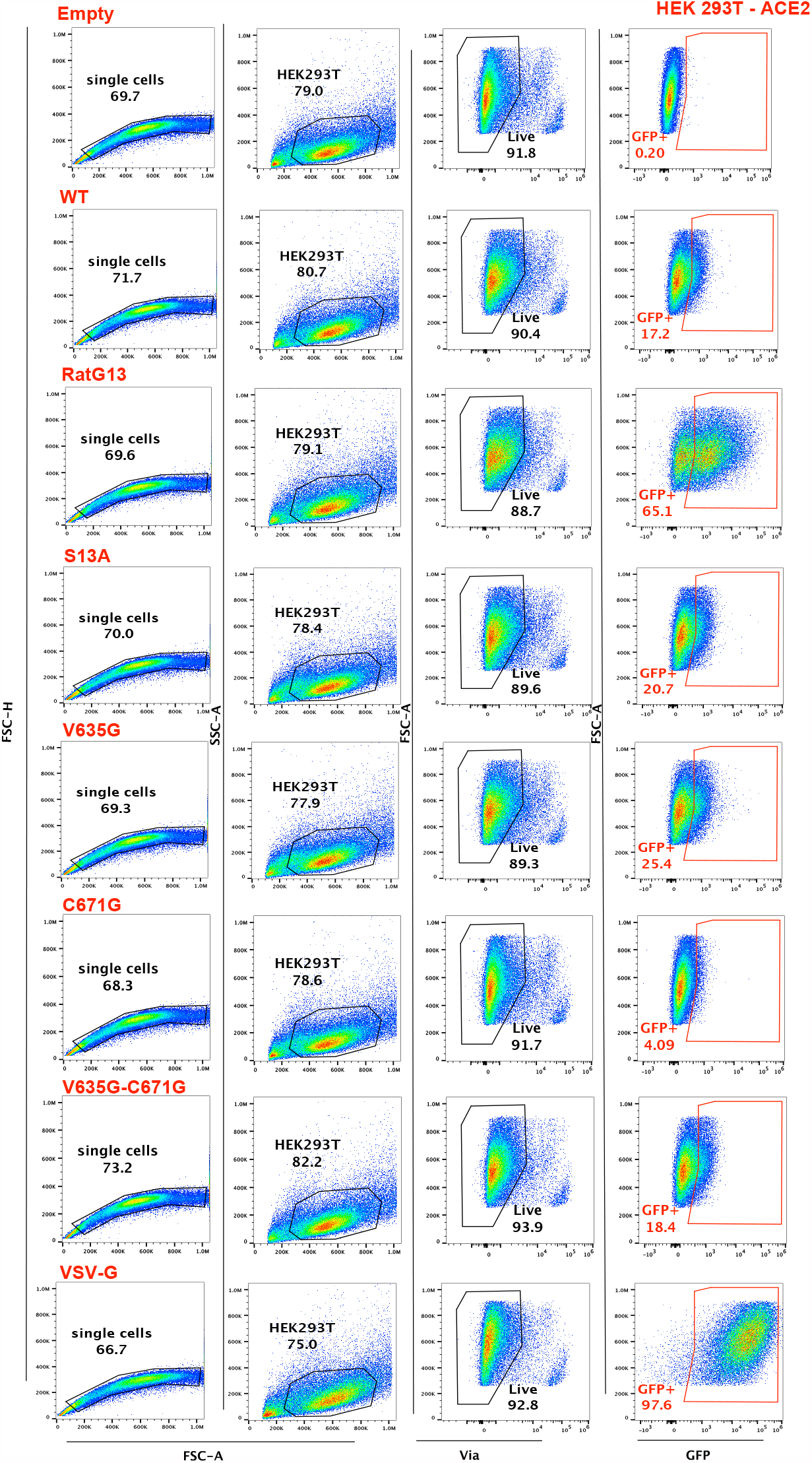
Scatter plots illustrating representative data and gating strategy for pseudovirus experiments performed in 293-Ace2 cells. 293-Ace2 cells were infected with spike-pseudotyped lentivectors expressing green fluorescent protein (GFP). Infection was quantified by measuring the percentage of GFP+ cells two days post-infection by flow cytometry. Cells were gated on FSC-H/FSC-A to exclude doublets, and then on SSC-A/FSC-A for size and granularity. Exclusion of dead cells was applied based on labeling for the Via-APC-eFluor780 dye, and infection was then measured based on the percentage of GFP+ cells among live cells.

**Fig. S8.**
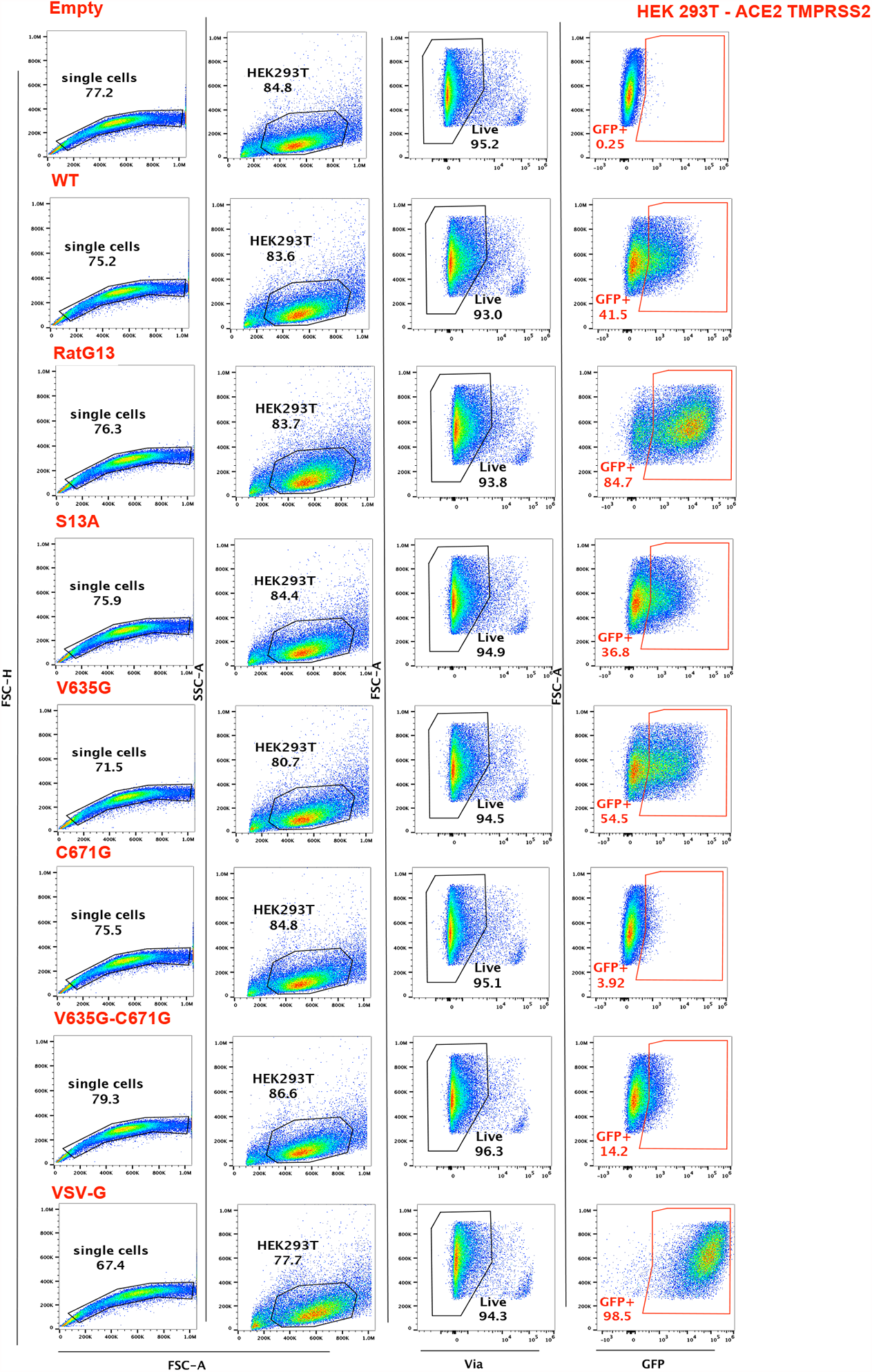
Scatter plots illustrating representative data and gating strategy for pseudovirus experiments performed in 293-Ace2-TMPRSS2 cells. 293-Ace2-TMPRSS2 cells were infected with spike-pseudotyped lentivectors expressing green fluorescent protein (GFP). Infection was quantified by measuring the percentage of GFP+ cells two days post-infection by flow cytometry. Cells were gated on FSC-H/FSC-A to exclude doublets, and then on SSC-A/FSC-A for size and granularity. Exclusion of dead cells was applied based on labeling for the Via-APC-eFluor780 dye, and infection was then measured based on the percentage of GFP+ cells among live cells.

**Fig. S9.**
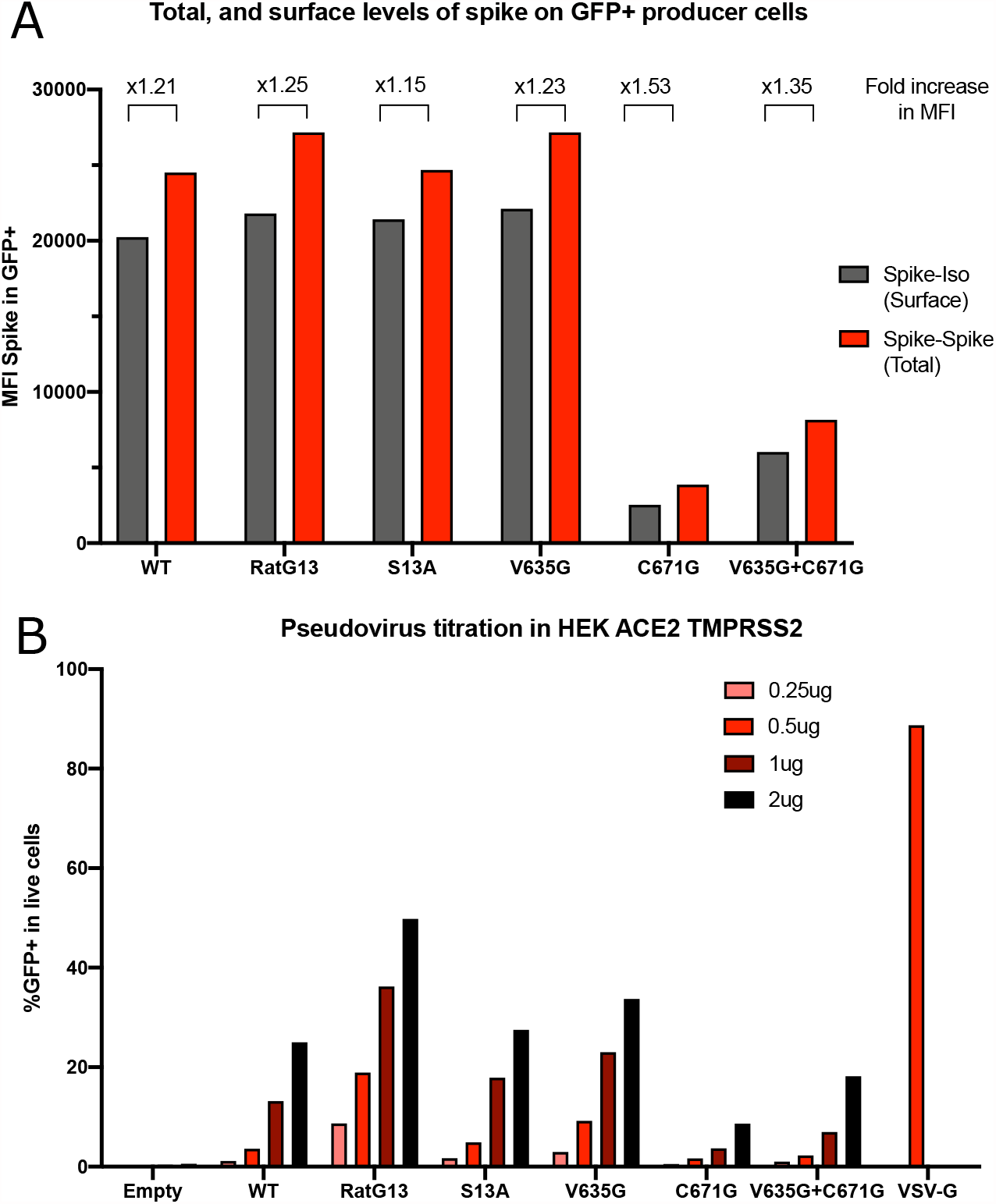
Extended characterisation of spike pseudotyped lentivectors. A) Mean fluorescence intensity plot examining levels of spike on the surface (spike-isotype control) and total levels of spike (surface and intracellular staining, spike-spike) in transfected (GFP+) HEK 293Tn producer cells (n = 1 biological replicate). Results indicate spike C671G single or V635G/C671G double mutants are present at lower levels in total and at the cell surface, suggesting a defect in stability rather than trafficking to the cell surface. Numbers over columns indicate the fold increase in MFI of total as compared to surface spike. B) Flow cytometry analysis of HEK-ACE2-TMPRSS2 cells transduced with a dilution series of the different spike pseudotyped lentivectors confirms the observations in Figure 3 remain consistent when input virus is scaled (n = 1 biological replicate). The amount of lentivector used is reported in p24 Gag protein equivalent (legend). The percentage of transduced cells (GFP+) is used to monitor infectivity.

**Fig. S10.**
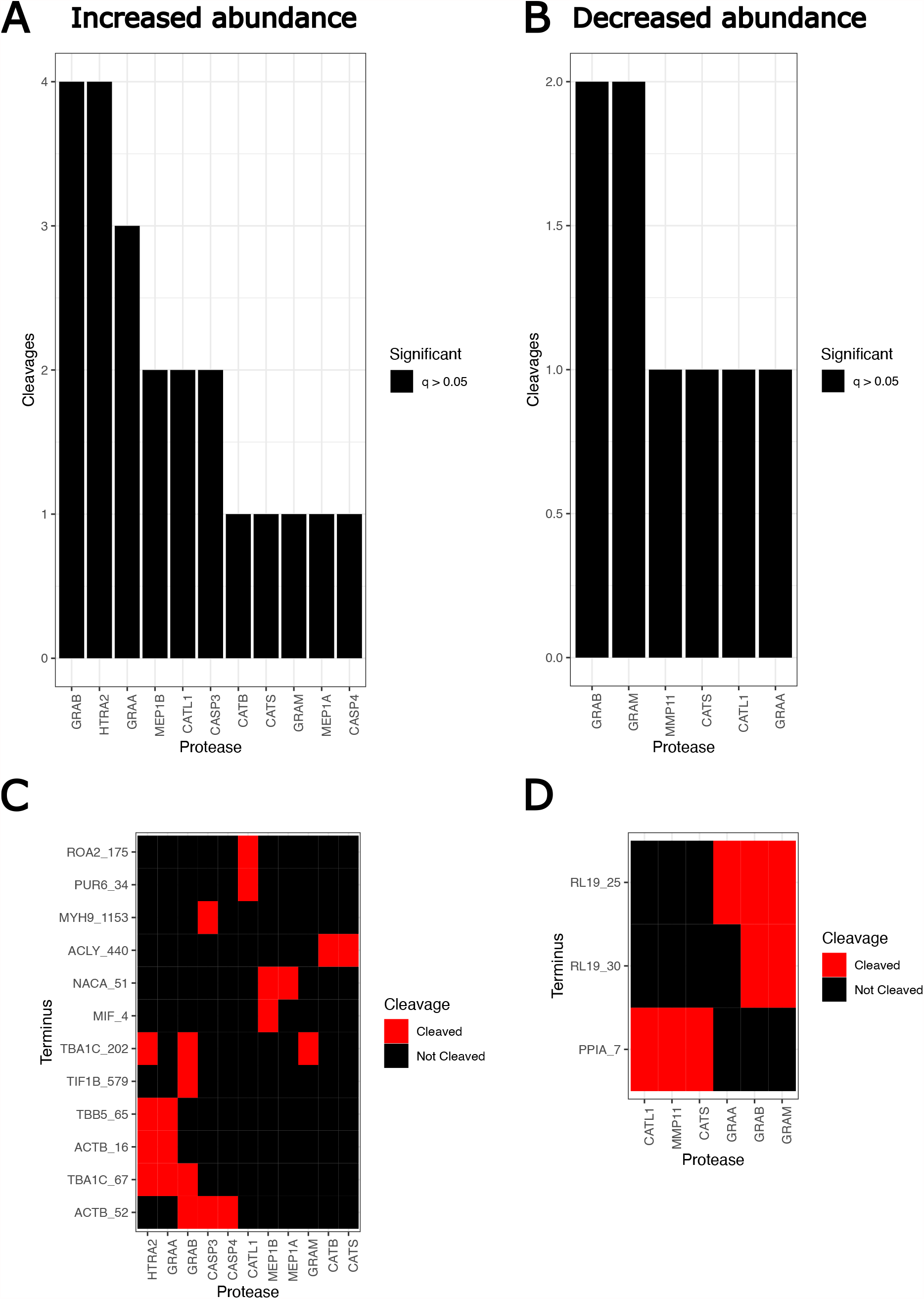
TopFIND analysis of cellular neo-N-termini significantly increased or decreased at 24h post-infection relative to mock (n = 3 biological replicates, Linma, Storeys Q-value <= 0.05). a) and b) show the number of neo-N-termini in each dataset associated with each causal protease. c) and d) show heatmaps matching individual neo-N-termini to proteases. Overall, TopFIND did not identify overrepresentation of substrates with these individual proteases (p <= 0.05, Adjusted Fisher’s exact test).

**Fig. S11.**
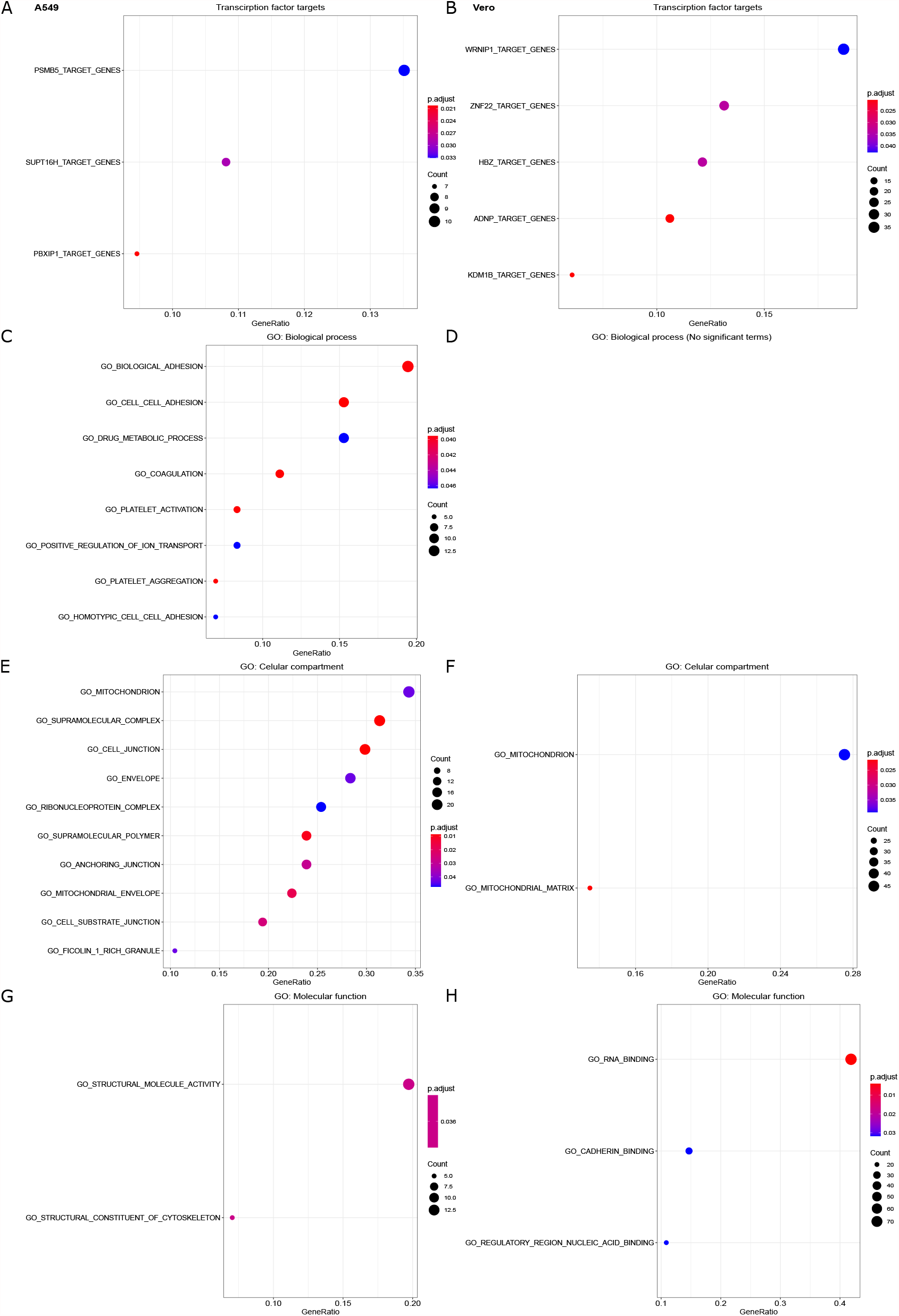
Differentially-expressed (Linma, Storeys Q-value adjusted P<=0.05) neo-N-termini from A549-Ace2 and Vero E6 cells at 24h post-infection compared to mock were analysed to determine functional enrichment. Here we show significantly-enriched A/B) transcription-factor targets, as well as gene ontology enrichment for C/D) Biological Processes, E/F) Cellular compartments G/H) Molecular functions.

**Fig. S12.**
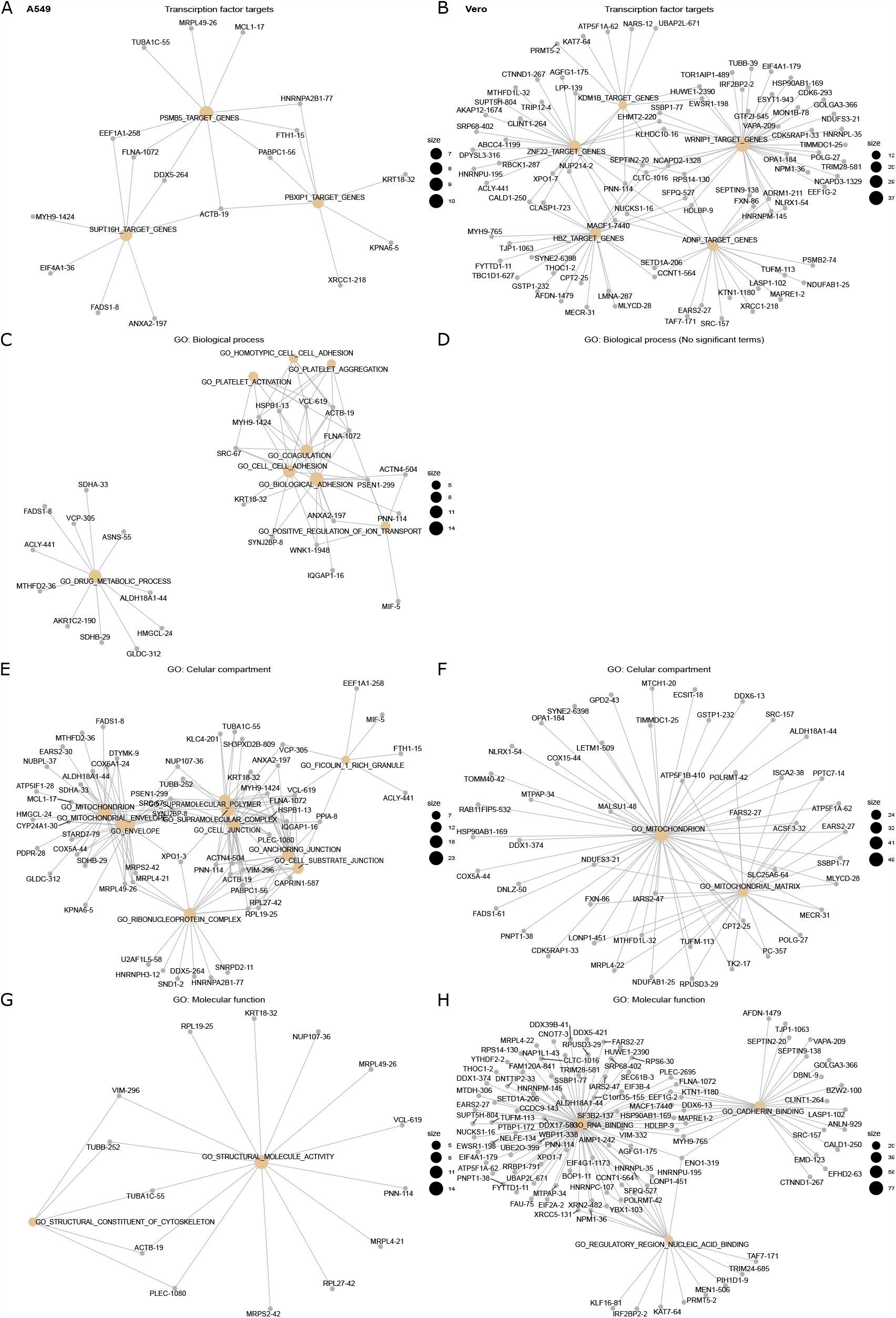
Differentially-expressed (Linma, Storeys Q-value adjusted P<=0.05) neo-N-termini from A549-Ace2 and Vero E6 cells at 24h post-infection compared to mock were analysed to determine functional enrichment. Here we highlight the networks of neo-N-termini associated with enriched A/B) transcription factor targets, as well as gene ontology enrichment for C/D) Biological Processes, E/F) Cellular compartments G/H) Molecular. The neo-N-termini are labeled as gene name_cleavage site position.

**Fig. S13.**
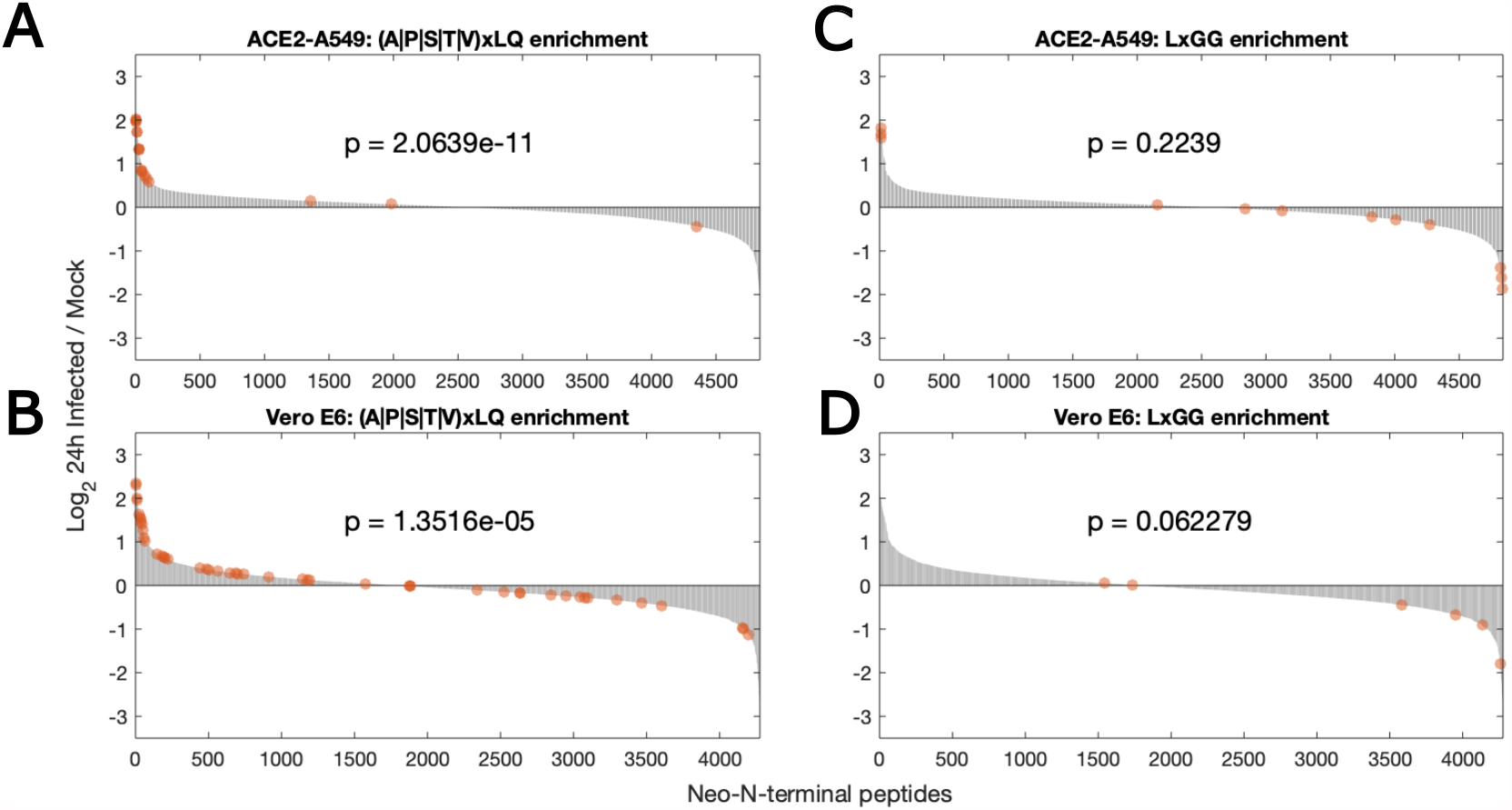
a) and b) Distribution of neo-N-termini consistent with the Mpro consensus motif in A549-Ace2 and Vero E6 cells respectively. c) and d) Distribution of neo-N-termini consistent with the PLP consensus motif in A549-Ace2 and Vero E6 cells respectively. Distributions cover all three biological replicates. Enrichment was determined by two-tailed Kolmogorov-Smirnov test.

**Fig. S14.**
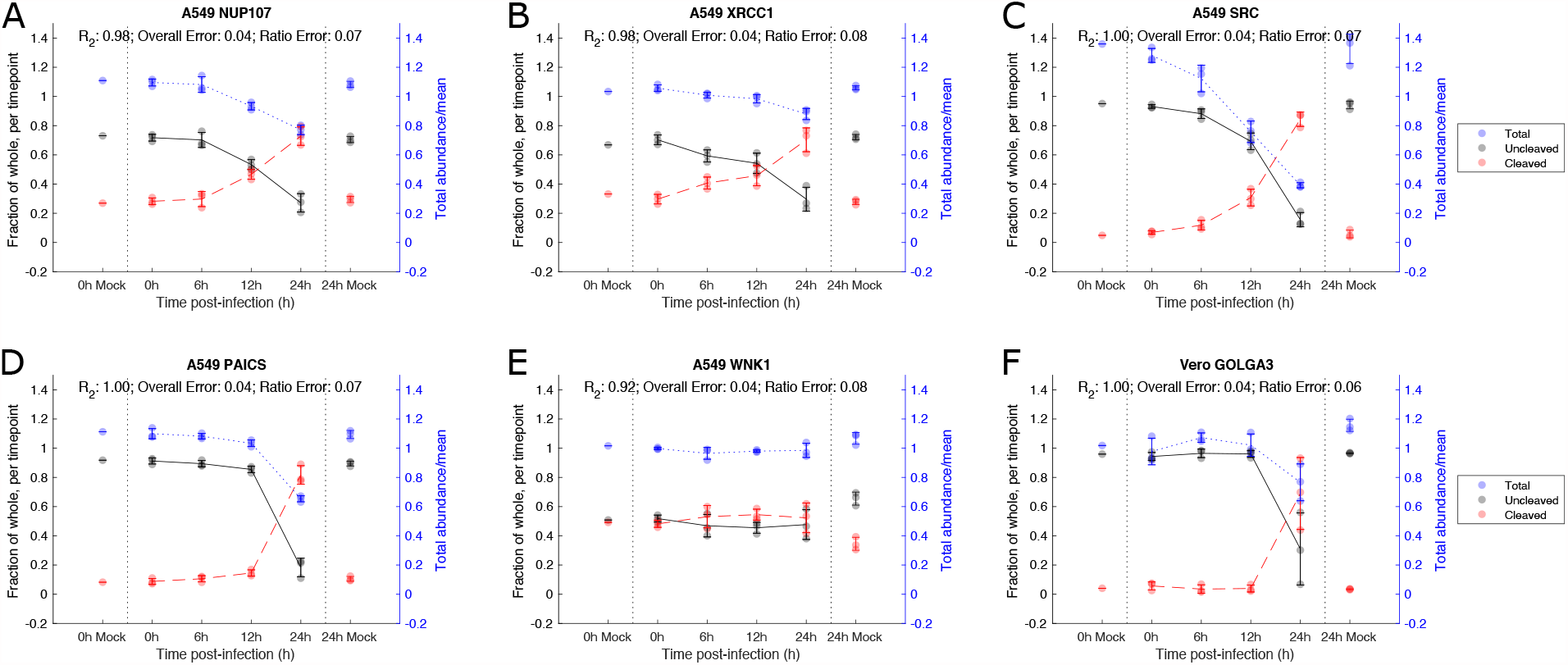
Cleavage stoichiometry for cellular proteins cleaved during viral infection was inferred using the HIquant approach (Malioutov et al. 2019) . The abundance of cleaved and uncleaved forms of each substrate are calculated on a per-timepoint basis. The total abundance (blue, right axis) is calculated by dividing the total abundance (cleaved and uncleaved) of protein at that timepoint by the mean total abundance accross all timepoints). n = 3 biological replicates.

**Fig. S15.**
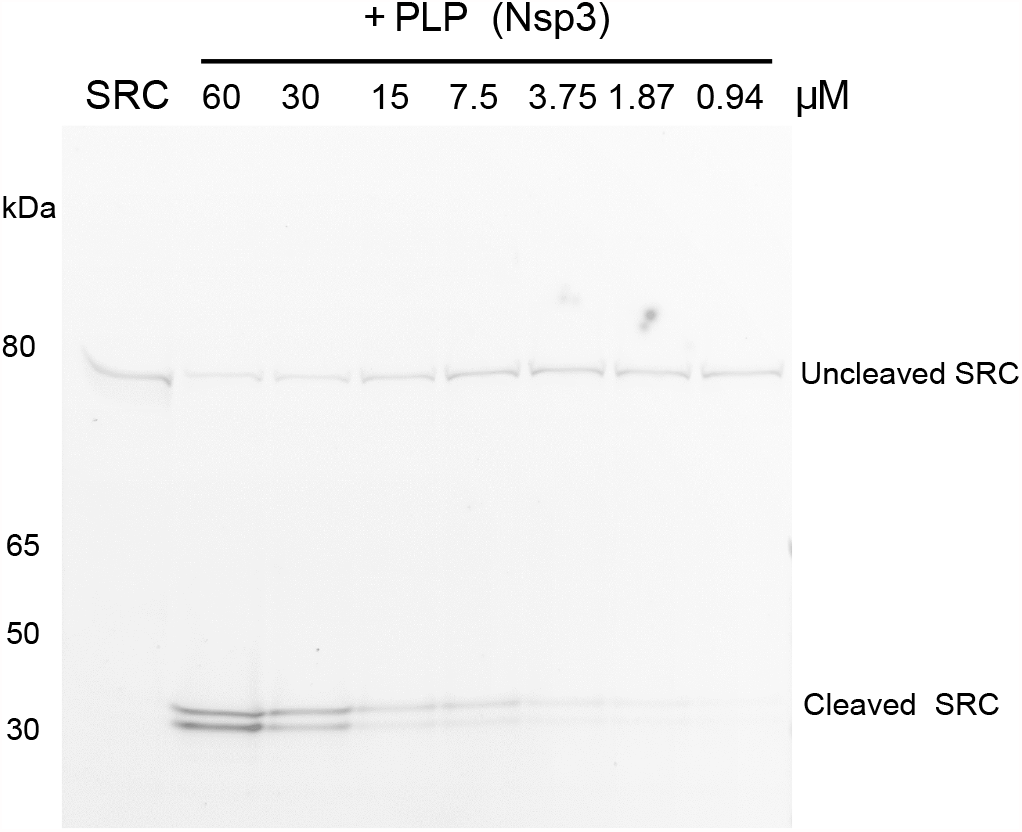
*In vitro*-translated N-terminally GFP-tagged SRC incubated with the indicated concentrations of SARS-CoV-2 PLP shows dose-dependent cleavage of SRC by PLP.

**Fig. S16.**
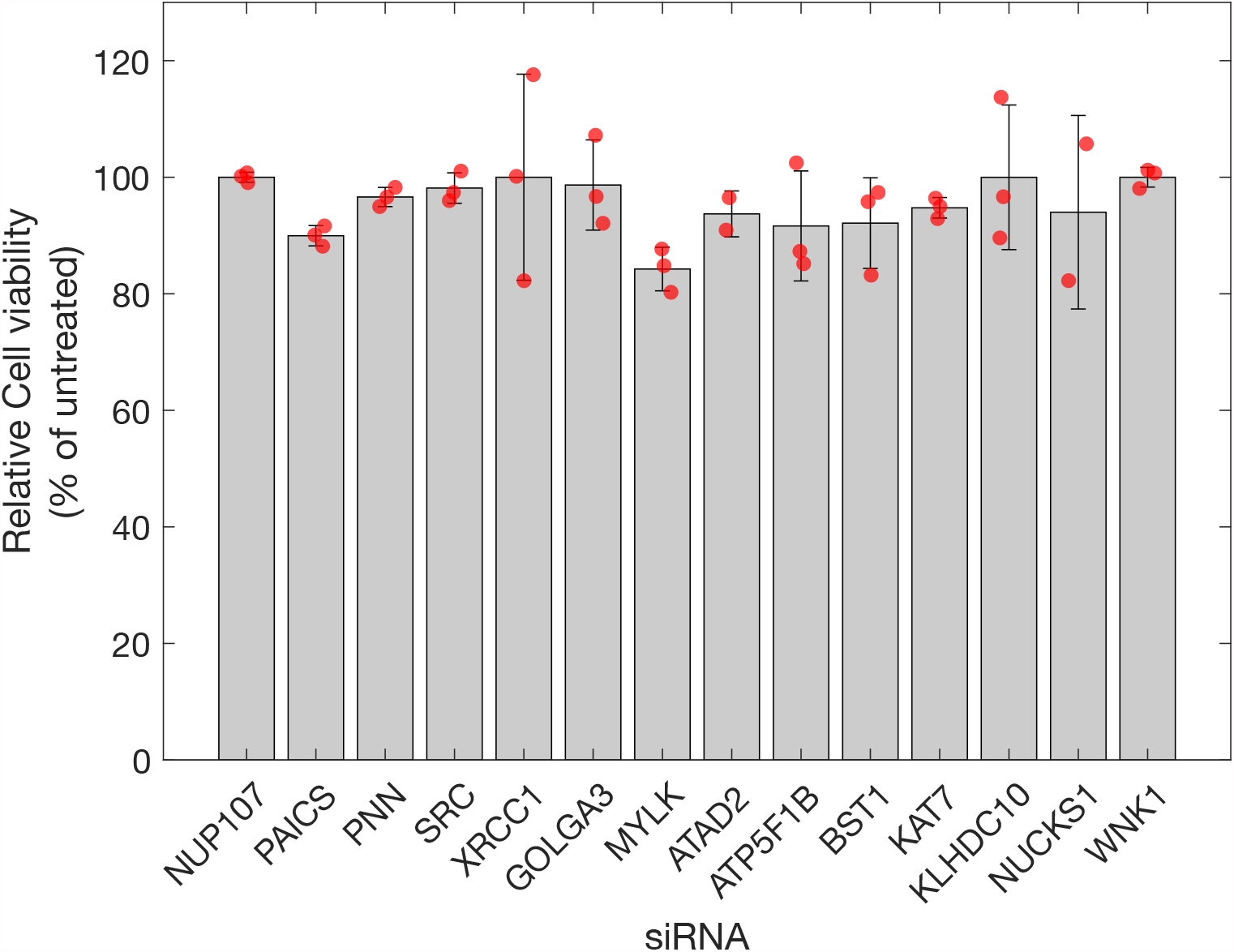
Cell viability of siRNA-treated A549-Ace2 cells. Cell viability was assessed by alamarBlueAlamar blue staining and compared to untreated control cells, and a 20% ethanol-lysed control. Error bars represent standard deviation from 3 biological replicates. Red markers indicate individual datapoints.

**Fig. S17.**
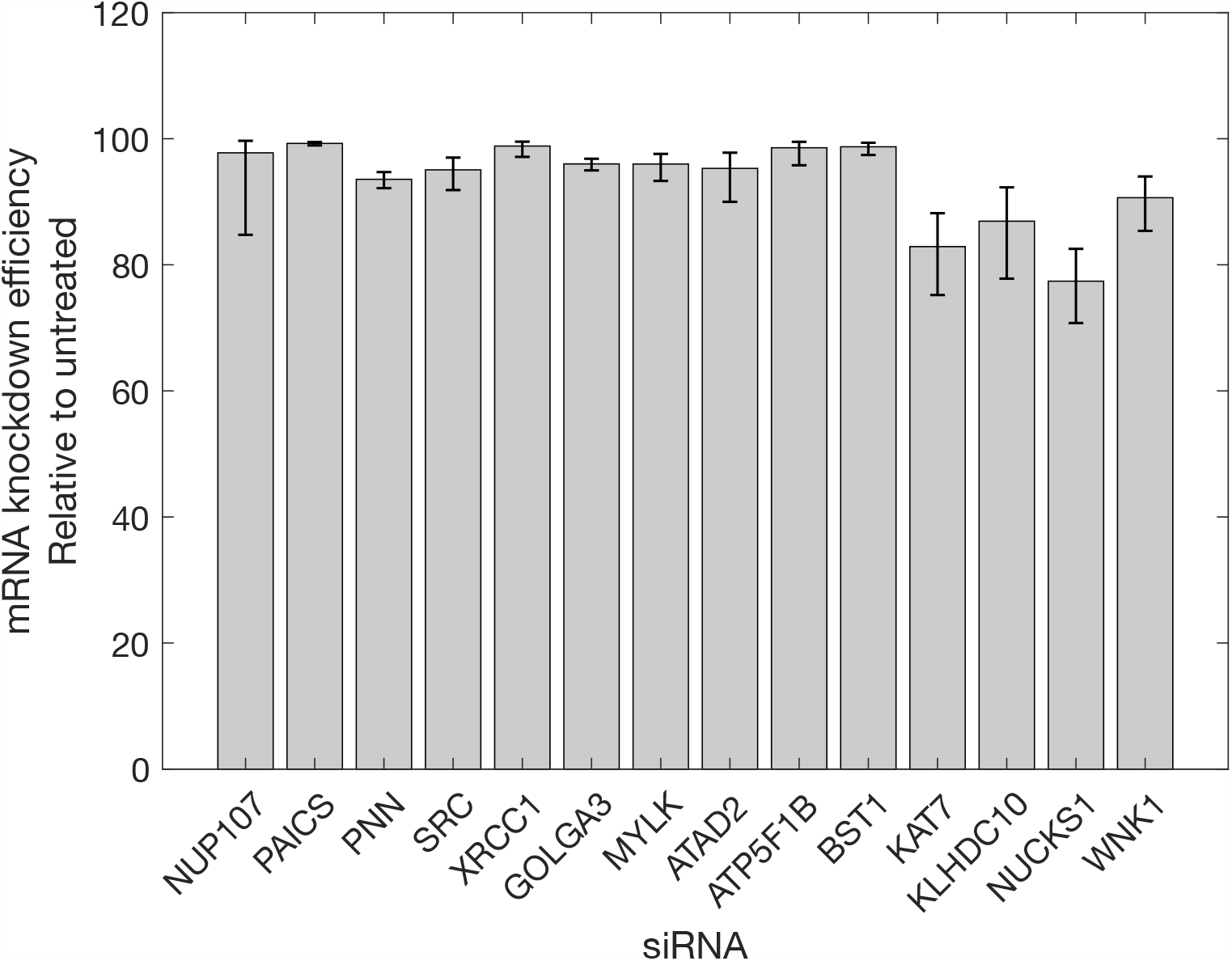
mRNA knockdown efficiency for SARS-CoV-2 protease substrates in siRNA-treated A549-Ace2 cells. Knockdown efficiency was calculated by qRT-PCR compared to a untreated control by the 2^*−*ΔΔ*Ct*^ method. Error bars represent standard deviation from a minimum of 3 biological replicates.

**Fig. S18.**
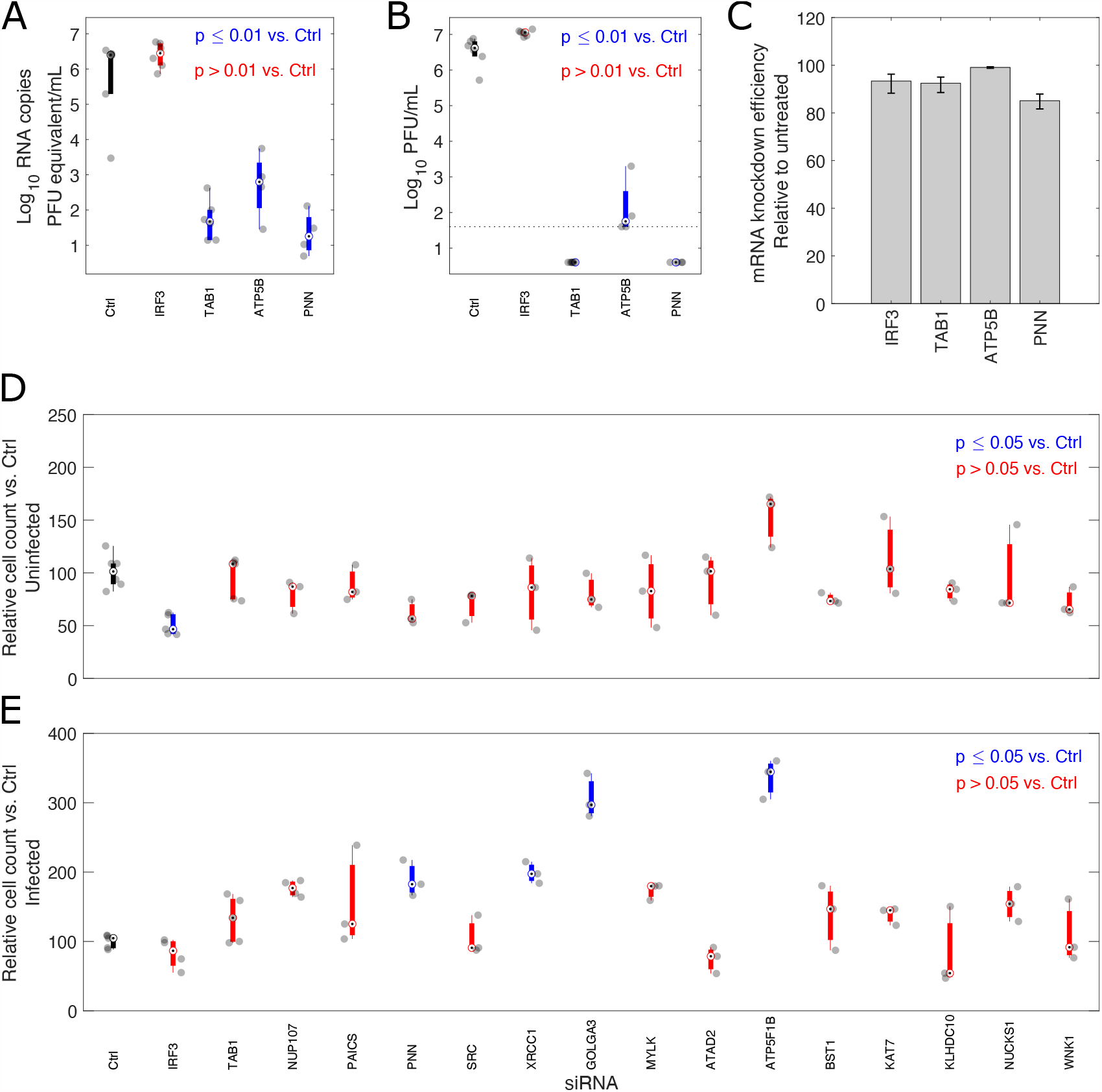
siRNA validation on previously identified SARS-CoV-2 protease substrates, including the antiviral substrate IRF3. A) SARS-CoV-2 RNA copies, B) Viral titres (PFU = plaque forming units) and C) KO efficiency relative to a scrambled siRNA control calculated by the 2^*−*ΔΔ*Ct*^ method. Relative cell numbers as determined by counting Hoechst 33258-stained nuclei in D) Mock and E) Infected cells, 72h post-infection/mock infection. For boxplots, circles with dots represent the median, the thick line the interquartile range, and whiskers extend to the furthest non-outlier datapoints. n *≥* 3 biological replicates. In panel C, the bars represent mean +/-standard deviation. Significance was determined by One-way ANOVA, using Tukey’s correction for multiple comparisons.

**Fig. S19.**
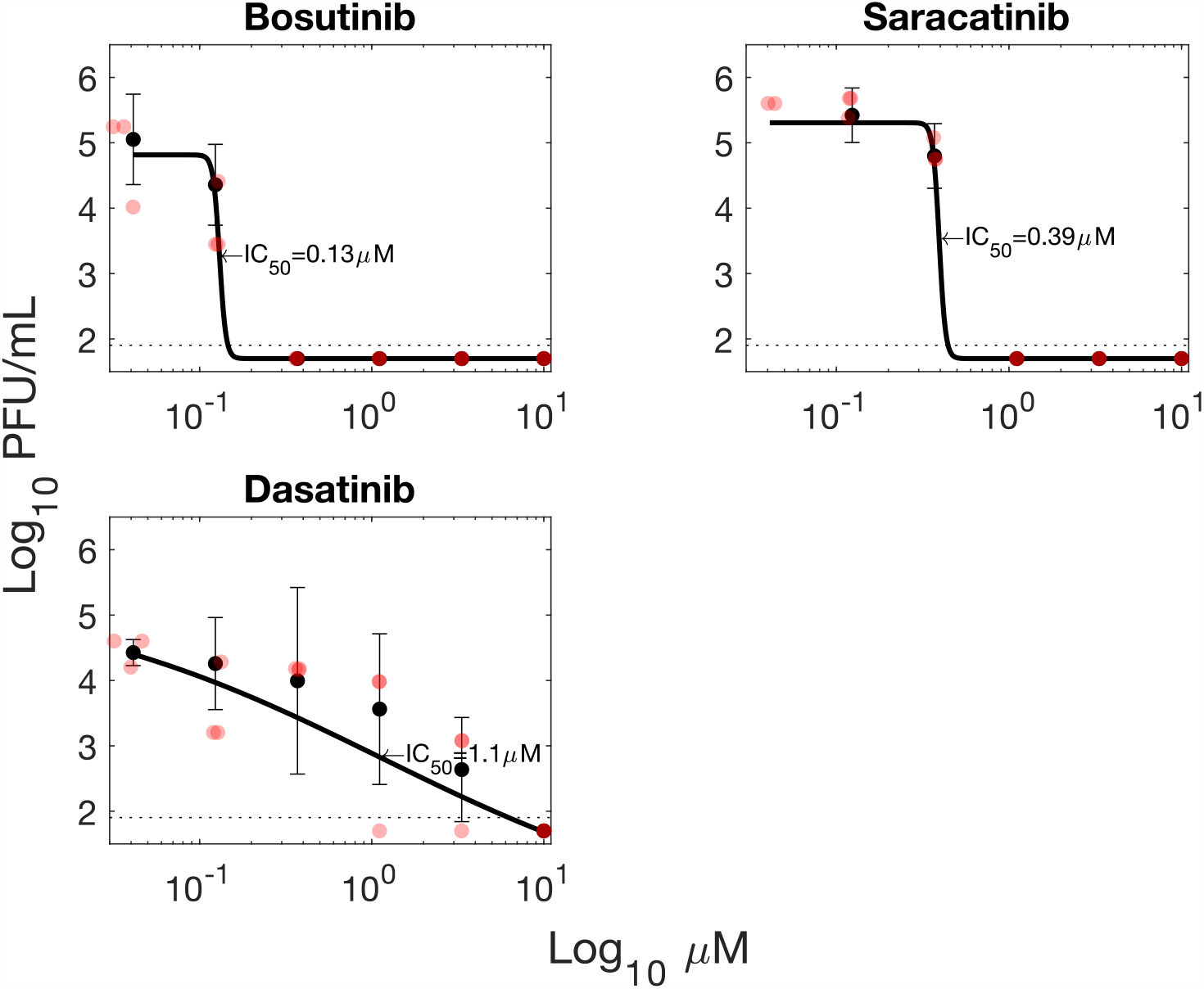
Additional Inhibitors targeting SRC kinase reduce SARS-CoV-2 titres in A549-Ace2 cells. Error bars represent standard deviation from 3 biological replicates. Red circles indicate individual datapoints.

**Fig. S20.**
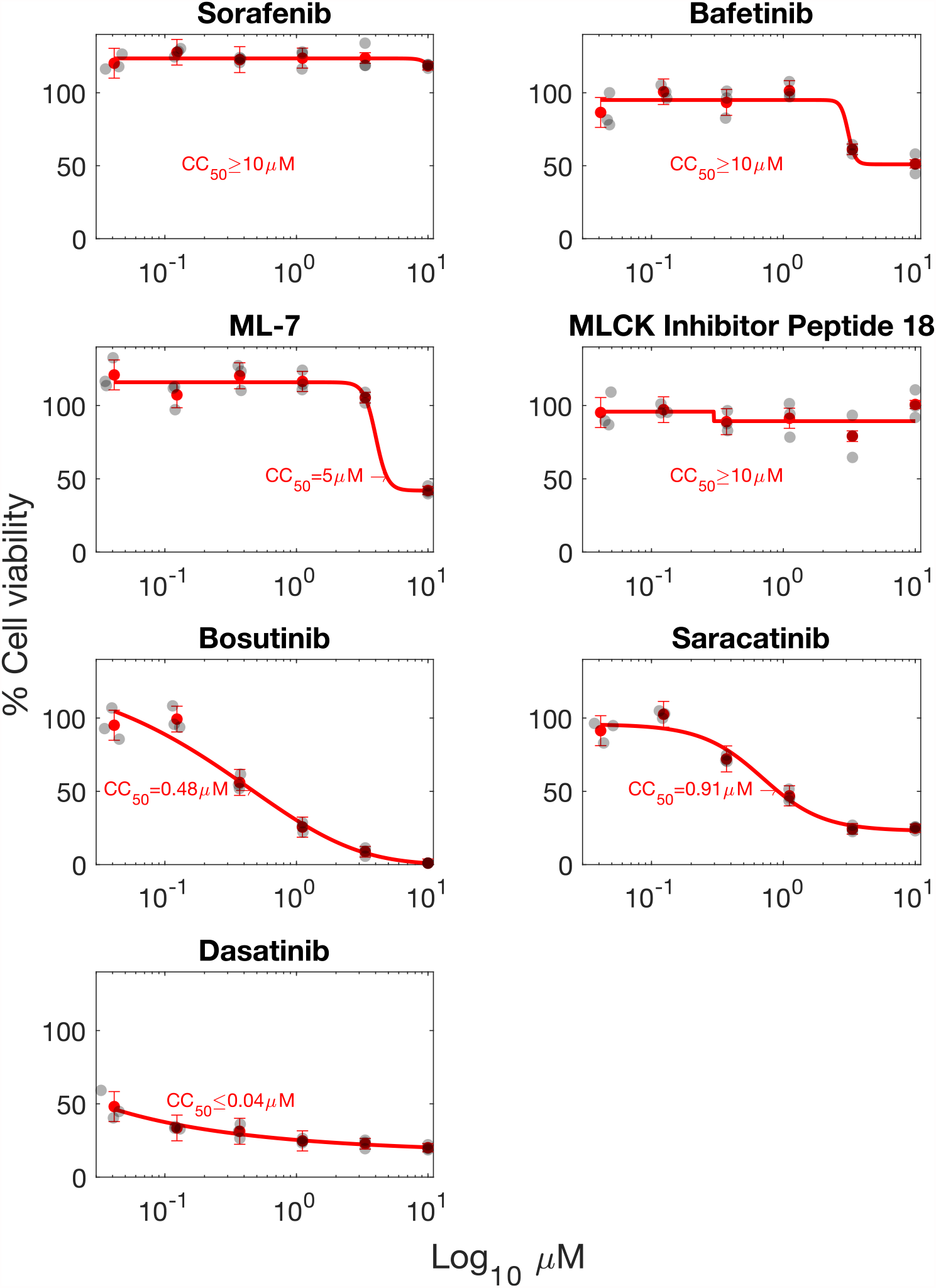
Cell viability and *CC*50 calculations for inhibitor-treated A549-Ace2 cells. Cell viability was assessed by Celltiter Glo staining and compared to untreated control cells, and a 20% ethanol-lysed control. Line represents best fit. Error bars represent standard deviation from 3 biological replicates. Black markers indicate individual data points.

